# Combination of Antiviral Drugs to Inhibit SARS-CoV-2 Polymerase and Exonuclease as Potential COVID-19 Therapeutics

**DOI:** 10.1101/2021.07.21.453274

**Authors:** Xuanting Wang, Carolina Q. Sacramento, Steffen Jockusch, Otávio Augusto Chaves, Chuanjuan Tao, Natalia Fintelman-Rodrigues, Minchen Chien, Jairo R. Temerozo, Xiaoxu Li, Shiv Kumar, Wei Xie, Dinshaw J. Patel, Cindy Meyer, Aitor Garzia, Thomas Tuschl, Patrícia T. Bozza, James J. Russo, Thiago Moreno L. Souza, Jingyue Ju

## Abstract

SARS-CoV-2 has an exonuclease-based proofreader, which removes nucleotide inhibitors such as Remdesivir that are incorporated into the viral RNA during replication, reducing the efficacy of these drugs for treating COVID-19. Combinations of inhibitors of both the viral RNA-dependent RNA polymerase and the exonuclease could overcome this deficiency. Here we report the identification of hepatitis C virus NS5A inhibitors Pibrentasvir and Ombitasvir as SARS-CoV-2 exonuclease inhibitors. In the presence of Pibrentasvir, RNAs terminated with the active forms of the prodrugs Sofosbuvir, Remdesivir, Favipiravir, Molnupiravir and AT-527 were largely protected from excision by the exonuclease, while in the absence of Pibrentasvir, there was rapid excision. Due to its unique structure, Tenofovir-terminated RNA was highly resistant to exonuclease excision even in the absence of Pibrentasvir. Viral cell culture studies also demonstrate significant synergy using this combination strategy. This study supports the use of combination drugs that inhibit both the SARS-CoV-2 polymerase and exonuclease for effective COVID-19 treatment.

## Introduction

Severe acute respiratory syndrome coronavirus 2 (SARS-CoV-2), which is responsible for COVID-19, is a positive-sense single-stranded RNA virus^1, 2^. Thus it requires an RNA-dependent RNA polymerase (RdRp) to replicate and transcribe its genome. Because of its large genome (∼30 kb) and error-prone RdRp, SARS-CoV-2 also possesses a 3’-5’ exonuclease for proofreading to maintain the integrity of the genome^3^. The replication complex of coronaviruses consists of several viral proteins, including the RdRp itself (nonstructural protein 12; nsp12) and its two accessory proteins (nsp7 and nsp8)^4–6^, and the exonuclease (nsp14) with its accessory protein (nsp10)^7^.

While a variety of drugs targeting many of the SARS-CoV-2 proteins essential for its infectious cycle have been evaluated, to date no effective antiviral strategy for COVID-19 exists. Remdesivir (RDV), a nucleotide analogue, is the only small molecule antiviral drug approved by the Food and Drug Administration (FDA) for COVID-19 treatment, although it was shown to have limited benefit in the World Health Organization’s Solidarity Trial. Lack of oral bioavailability and short half-life reduces RDV’s clinical effectiveness. With the appearance of new SARS-CoV-2 variants that pose a threat to public health and may escape the immune response to current vaccines, it is more critical than ever to develop antivirals that can treat COVID-19 effectively. We have investigated a library of nucleotide analogues targeting the SARS-CoV-2 RdRp, several of which have been FDA approved for other viral infections including HIV/AIDS and hepatitis B and C^8–11^. Of particular interest are those that are able to be incorporated into the replicated RNA by the viral RdRp where they halt or slow further replication. These include, among several others, nucleotide analogues such as Sofosbuvir (Fig. 1a), an immediate terminator of the polymerase reaction which has since entered COVID-19 clinical trials^12, 13^, Remdesivir (Fig. 1b), a delayed terminator^14, 15^, and Tenofovir, an obligate terminator (Fig. 1d)^10, 11^. Recently, we demonstrated that upon incorporation of the triphosphate forms of Sofosbuvir and Remdesivir into RNA by the SARS-CoV-2 RdRp complex, both nucleotide analogues were removed by the SARS-CoV-2 exonuclease complex, although Sofosbuvir was removed at a slower rate^16^. Currently, three other prodrugs related to nucleotides, Favipiravir (Fig. 1c), Molnupiravir and AT-527 (Fig. 1e, f), are undergoing clinical trials for COVID-19.

**Fig. 1.**
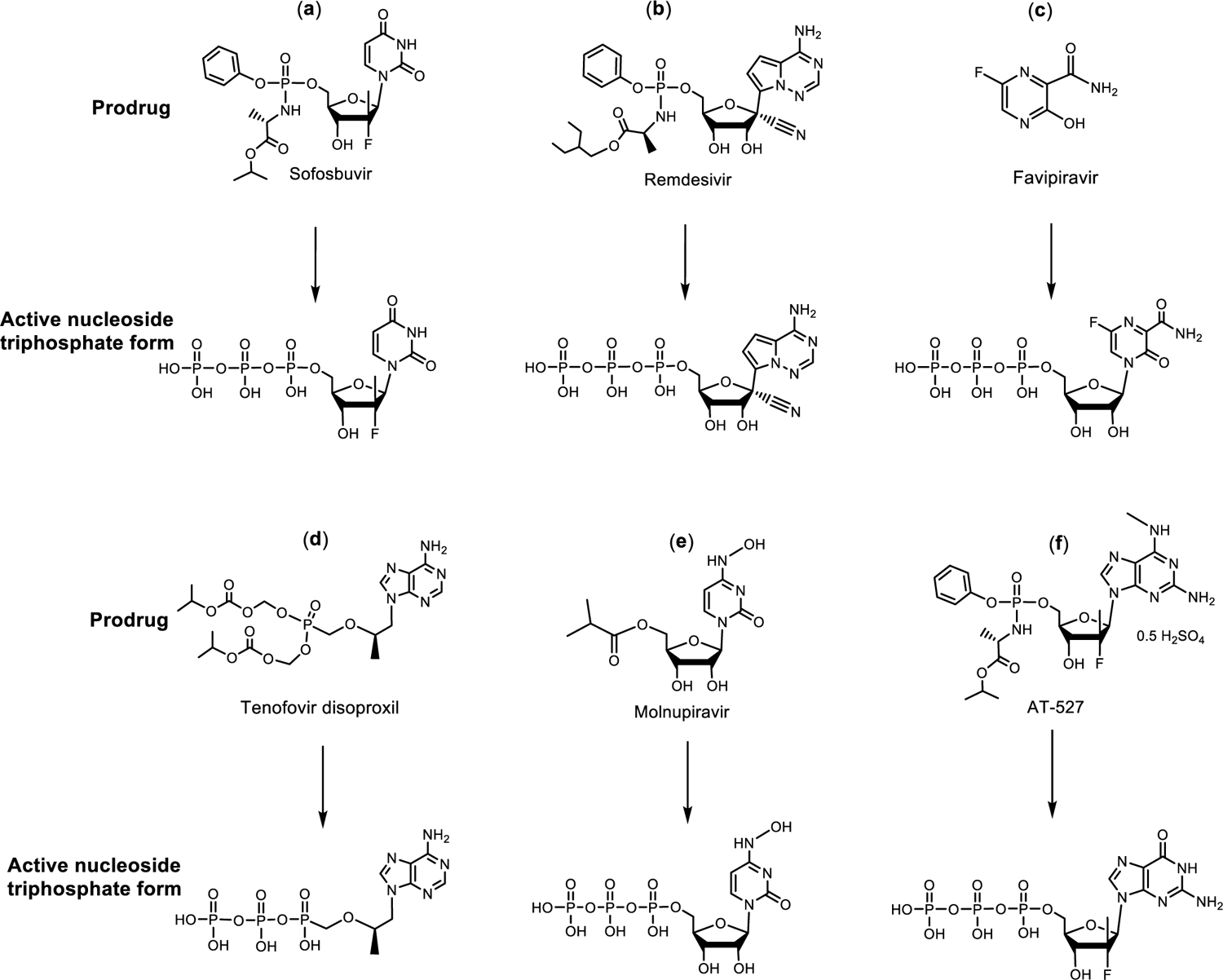
Structures of the prodrugs Sofosbuvir (a), Remdesivir (b), Favipiravir (c), Tenofovir disoproxil (d), Molnupiravir (e) and AT-527 (f) (top) and their respective active triphosphate forms Sofosbuvir-5’-triphosphate, Remdesivir-5’-triphosphate, Favipiravir-ribofuranosyl-5’-triphosphate (Favipiravir-RTP), Tenofovir diphosphate, N^4^-hydroxycytidine-5’-triphosphate (NHC-TP) and 2’-fluoro-2’-methyl guanosine-5’-triphosphate (AT-9010, G_fm_-TP) (bottom).

While nucleoside/nucleotide analogues show promising results against SARS-CoV-2, *in vitro* pharmacological parameters often exceed human plasma exposure. Most likely, nucleoside/nucleotide-based inhibitors are insufficient to block viral RNA replication, because they could be quickly excised by the SARS-CoV-2 exonuclease. Combination of RdRp inhibitors such as Sofosbuvir, Remdesivir and others listed in Fig. 1 with exonuclease inhibitors may provide a more effective strategy for blocking SARS-CoV-2 RNA replication. We previously described the need to overcome the exonuclease proofreading function in order to completely inhibit SARS-CoV-2 replication in early 2020.^10^ An *in silico* study has proposed combining RdRp and exonuclease inhibitors as a strategy to combat COVID-19^17^; however, no successful demonstration of such a combined drug strategy has been reported. Hepatitis C virus (HCV) and SARS-CoV-2 are both positive sense single stranded RNA viruses. Viruses with small genomes that encode relatively few proteins, such as HCV, must possess multifunctional proteins^18^. HCV NS5A is a good example because it binds lipids, Zn^++^ and RNA, as well as participates in phosphorylation and cell signaling events^18^. The SARS-CoV-2 genome is 3-fold larger than that of HCV, and the analogous NS5A functions are likely distributed among proteins nsp1 to nsp16 of the coronavirus^19^. We have previously shown that the NS5A inhibitors Daclatasvir and Velpatasvir inhibit both the SARS-CoV-2 polymerase and exonuclease activities^20, 21^. The combination of Velpatasvir with Remdesivir was shown to have a synergistic effect relative to Remdesivir alone in inhibiting SARS-CoV-2 replication^22^. Here we further examined whether the NS5A inhibitors Pibrentasvir, Ombitasvir and Daclatasvir (structures shown in Fig. S-1), possess anti-exonuclease activity and could synergize with RdRp inhibitors in virus replication assays. We discovered that Pibrentasvir and Ombitasvir have the highest exonuclease inhibitory activity among all the NS5A inhibitors we have tested. As an example, we show here that Pibrentasvir reduced the removal by exonuclease of immediate and delayed RNA chain terminators incorporated into the RNA. Consequently, Pibrentasvir and Ombitasvir synergistically improved the *in vitro* pharmacological parameters of several RdRp inhibitors, such as Sofosbuvir, Remdesivir, Tenofovir and Favipiravir. Since these RdRp inhibitors are clinically approved antivirals, candidate combinations of the polymerase and exonuclease inhibitors identified here could be prioritized for advancement to COVID-19 clinical trials.

## Results

### Molecular docking study of HCV NS5A inhibitors with SARS-CoV-2 exonuclease nsp14

We performed a molecular docking study to investigate whether the HCV NS5A inhibitors (Fig. S-1) interact with the SARS-CoV-2 exonuclease. Since a 3D structure for SARS-CoV-2 nsp14 was not yet available, we built a protein model based on the ortholog in SARS-CoV^3^. Superposition of the 3D structures of these two enzymes indicates that the main domains and the catalytic amino acid residues of the exonuclease active sites are structurally similar (Fig. S-2). The clinically approved HCV NS5A inhibitors are bound to the SARS-CoV-2 nsp14 exonuclease active site with comparable docking scores (Fig. 2). The exonuclease activity requires the proper coordination of the Mg^++^ ion with amino acid residues Asp-90, Glu-92, Glu-191 and Asp-273 as well as the RNA substrate. NS5A inhibitors interfere with the coordination among the Mg^++^ ion, the amino acids in the nsp14 active site and the RNA 3’ terminus, likely preventing nucleotide excision from the RNA (Fig. 2).

**Fig. 2.**
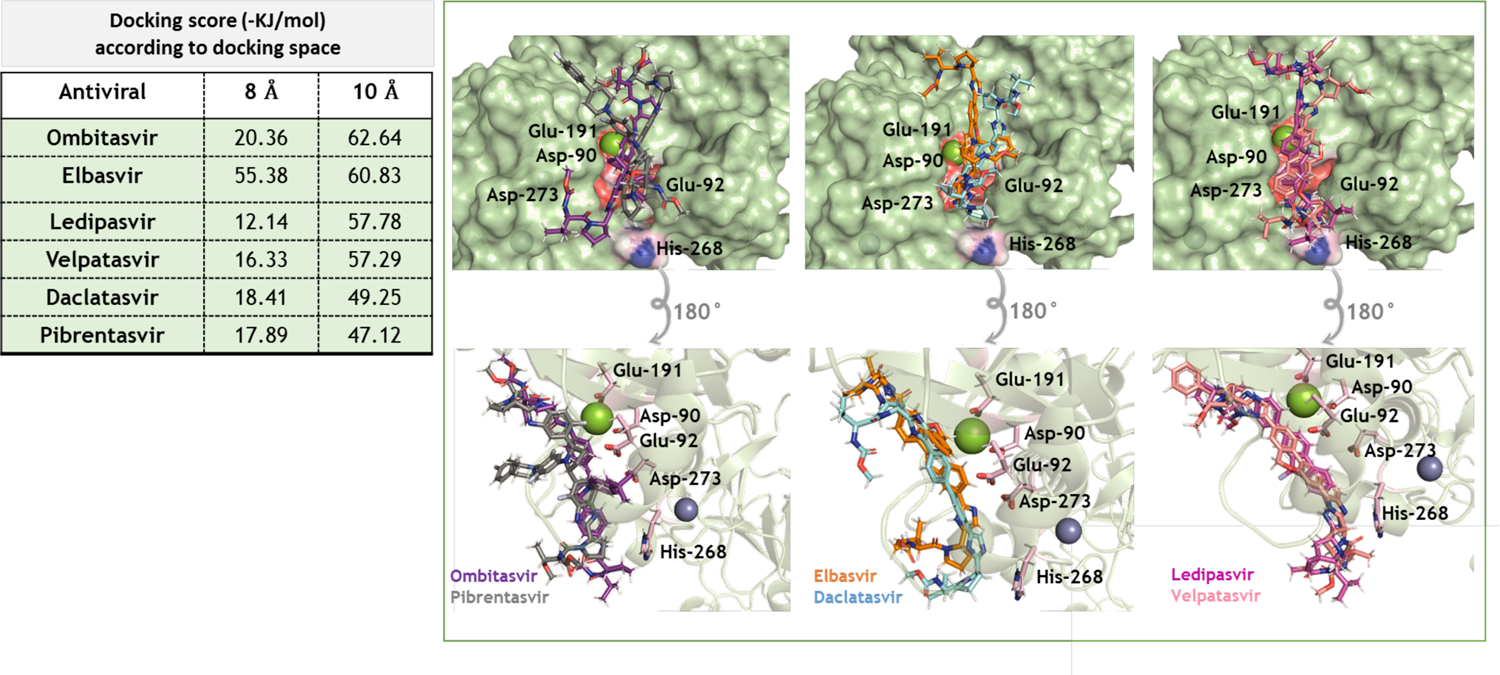
3D representation of the best docking poses for HCV NS5A inhibitors in the nsp14 exonuclease active site. The NS5A inhibitors were built and minimized in terms of energy by Density Functional Theory (DFT). Docking was performed using GOLD 2020.2 software with *ChemPLP* as scoring function. Because of the diversity in NS5A inhibitors’ molecular weights, they were allowed to dock within 8 or 10 Å spheres in the SARS-CoV-2 nsp14 exonuclease active site, constructed in Fig. S-2. The NS5A and catalytic amino acid residues are in stick representation. Mg^++^ and Zn^++^ are represented as green and indigo blue spheres, respectively.

### Inhibition of SARS-CoV-2 exonuclease activity by HCV NS5A inhibitors to prevent the excision of immediate and delayed terminators from RNA

Using the molecular assays that we have developed previously^16^, we evaluated if HCV NS5A inhibitors could indeed inhibit SARS-CoV-2 exonuclease activity. We identified Pibrentasvir and Ombitasvir as two strong inhibitors of the SARS-CoV-2 exonuclease. As a proof of principle, we used Pibrentasvir to demonstrate whether such drugs can prevent excision of nucleotide analogues from RNAs terminated with the active forms of prodrugs such as Remdesivir, Sofosbuvir, Tenofovir, Favipiravir, Molnupiravir and AT-527.

Pibrentasvir and Ombitasvir inhibit the SARS-CoV-2 exonuclease in a concentration dependent manner, as shown in Fig. 3. RNA (sequence shown at the top of the figure) was incubated with a SARS-CoV-2 pre-assembled exonuclease complex (nsp14/nsp10) at 37 °C for 15 min in the absence (b) and presence of varying amounts of Pibrentasvir (c-e). The RNA (a) and the products of the exonuclease reaction (b-e) were analyzed by MALDI-TOF MS. The peak at 8160 Da corresponds to the intact RNA. In the absence of Pibrentasvir, exonuclease activity caused nucleotide cleavage from the 3’-end of the RNA as shown by the 7 lower molecular weight fragments corresponding to cleavage of 1-7 nucleotides (b), with only ∼7% intact RNA remaining (peak at 8167 Da). Pibrentasvir (1, 10 and 20 µM) reduced exonuclease activity in a concentration dependent manner as shown by the reduced intensities of the fragmentation peaks and more prominent intact RNA peak (c-e). Ombitasvir displayed similar efficiency of exonuclease activity inhibition (Fig. 3 f-i) as Pibrentasvir. Daclatasvir also exhibits inhibition of exonuclease activity, but at higher concentrations than either Pibrentasvir or Ombitasvir (Fig. S-3).

**Fig. 3.**
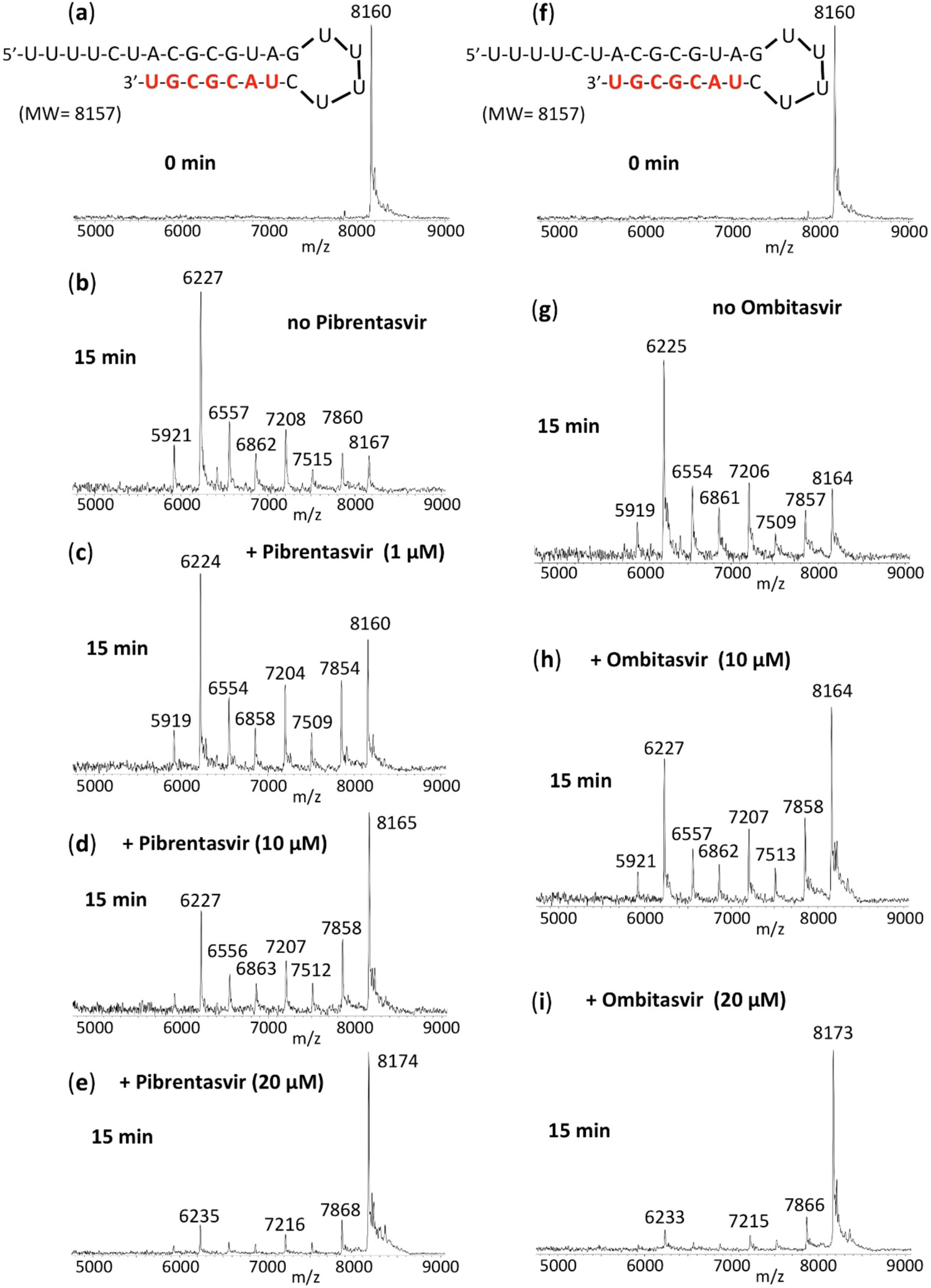
Inhibition of SARS-CoV-2 exonuclease activity by Pibrentasvir and Ombitasvir. A mixture of 500 nM RNA (sequence shown at the top of the figure) and 50 nM SARS-CoV-2 pre-assembled exonuclease complex (nsp14/nsp10) was incubated in buffer solution at 37 °C for 15 min in the absence (b, g) and presence of varying amounts of Pibrentasvir (c-e) and Ombitasvir (h, i). The RNA (a, f) and the products of the exonuclease reaction (b-e, g-i) were analyzed by MALDI-TOF MS. The signal intensity was normalized to the highest peak. The peak at 8160 Da corresponds to the intact RNA (8157 Da expected). In the absence of Pibrentasvir, exonuclease activity caused nucleotide cleavage from the 3’-end of the RNA as shown by the 7 lower molecular weight fragments corresponding to cleavage of 1-7 nucleotides (b). Under our experimental conditions, approximately 7% intact RNA remained as indicated by the peak at 8167 Da (b). With increasing amounts of Pibrentasvir, exonuclease activity was reduced as shown by the reduced intensities of the fragmentation peaks and increased intact RNA peak (c-e). Similar results were observed with Ombitasvir (f-i).

We next determined if RNA terminated with the active forms of prodrugs such as Sofosbuvir, Remdesivir, Tenofovir, Favipiravir, Molnupiravir and AT-527 were also less likely to be excised by SARS-CoV-2 exonuclease in the presence of Pibrentasvir. We generated Sofosbuvir and Remdesivir terminated RNA by single base extension of RNA template-loop-primers with the active triphosphate forms of Sofosbuvir and Remdesivir, respectively, using the replication complex assembled from SARS-CoV-2 nsp12 (the viral RdRp) and nsp7 and nsp8 proteins (RdRp cofactors)^11, 16^. The Sofosbuvir (S)-terminated RNA and Remdesivir (R)-terminated RNA (sequences shown at the top of Fig. 4) were incubated with the SARS-CoV-2 nsp14/nsp10 complex at 37 °C for 15 min in the absence (Fig. 4b, e) and presence of 20 µM Pibrentasvir (c, f). MALDI-TOF MS analysis of the intact RNAs (a, d) and the products of the exonuclease reactions (b, c, e, f) was performed. In the absence of Pibrentasvir, exonuclease activity caused cleavage of 1-7 nucleotides from the 3’-end of the RNA as shown by the lower molecular weight fragments (b, e). When 20 µM Pibrentasvir was added, exonuclease activity was substantially inhibited as shown by the reduced intensities of the fragmentation peaks and increased peak heights of the intact RNAs (c, f). Thus, the SARS-CoV-2 exonuclease activity is significantly inhibited by Pibrentasvir to prevent nucleotide excision from both Sofosbuvir and Remdesivir terminated RNA. Similarly, Pibrentasvir substantially reduced excision of the nucleotide analogues derived from Favipiravir (Fig. S-4), Molnupiravir (NHC, N^4^-hydroxycytidine, Fig. S-5) and AT-527 (Fig. S-6) from the 3’ terminus of RNA. In terms of base pairing, Favipiravir-RTP acts like a G or A^23^, NHC-TP behaves like a C or U^24^ and AT-9010 (the active triphosphate form of AT-527) is a G analogue^25^. As expected, Favipiravir-RTP was incorporated into RNA opposite C or U in the template strand (Fig. S-4), NHC-TP was incorporated opposite A or G (Fig. S-5), and AT-9010 was incorporated opposite C (Fig. S-6) in the template by the RdRp complex (nsp12/nsp7/nsp8). In all cases, the incorporated nucleotide analogue along with the adjacent 6-7 nucleotides were efficiently removed following incubation with the exonuclease complex (nsp14/nsp10) (Fig. S-4b, e; S-5e, h; S-6e). However, in the presence of 20 µM Pibrentasvir, this exonuclease activity was largely abrogated (Fig. S-4c, f; S-5f, i; S-6f).

**Fig. 4.**
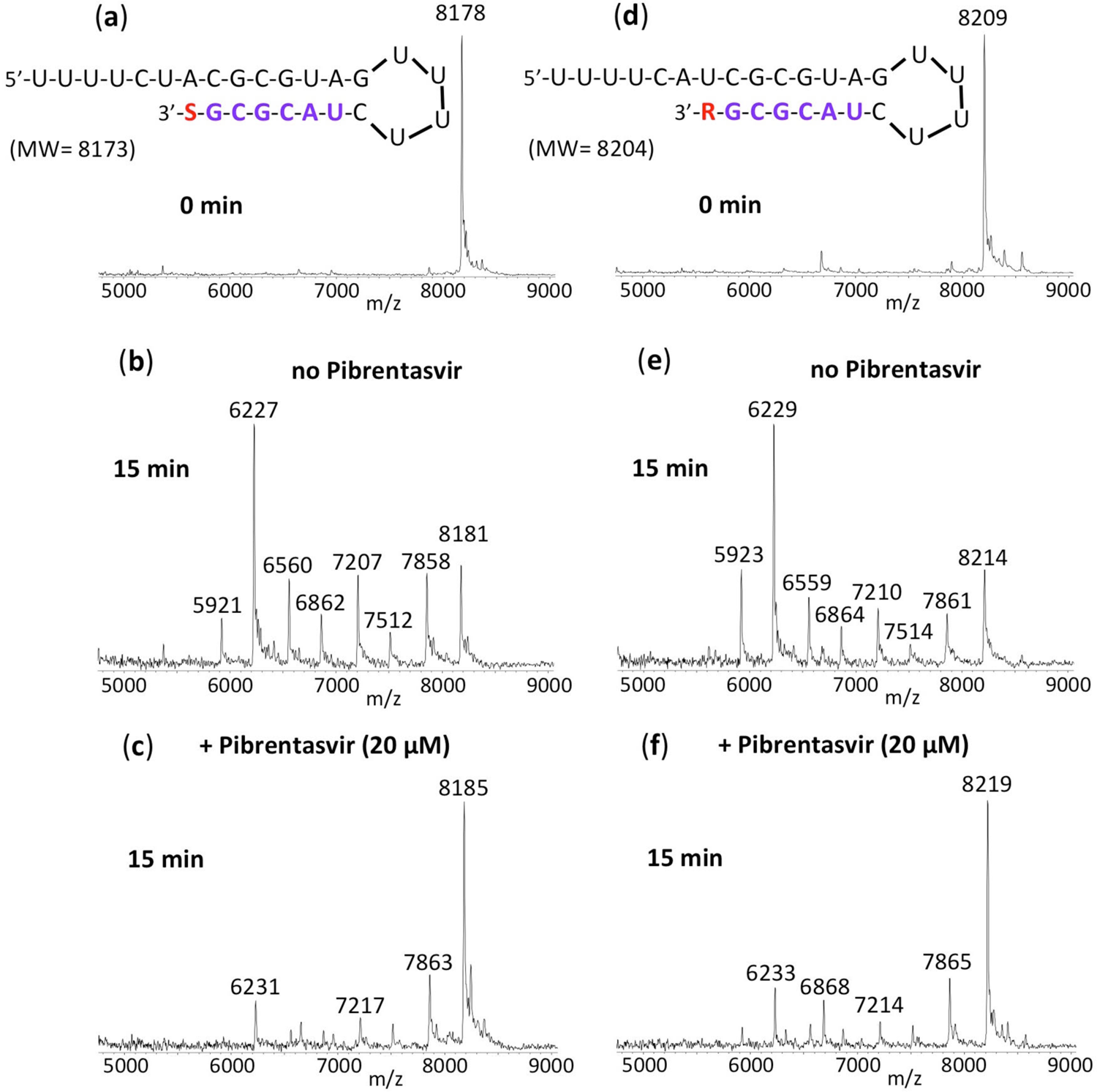
Inhibition of SARS-CoV-2 exonuclease activity by Pibrentasvir for Sofosbuvir (S) and Remdesivir (R) terminated RNA. A mixture of 500 nM RNAs (sequences shown at the top of the figure) and 50 nM SARS-CoV-2 pre-assembled exonuclease complex (nsp14/nsp10) were incubated in buffer solution at 37 °C for 15 min in the absence (b, e) and presence of 20 µM Pibrentasvir (c, f). The intact RNAs (a, d) and the products of the exonuclease reactions (b-f) were analyzed by MALDI-TOF MS. The signal intensity was normalized to the highest peak. In the absence of Pibrentasvir, exonuclease activity caused nucleotide cleavage from the 3’-end of the RNA as shown by the lower molecular weight fragments corresponding to cleavage of 1-7 nucleotides (b, e). When 20 µM Pibrentasvir was added, exonuclease activity was reduced as shown by the reduced intensities of the fragmentation peaks and increased intact RNA peaks (c, f).

The nucleotide inhibitor Remdesivir exhibits delayed termination during the polymerase reaction, which has been suggested as one of the possible mechanisms for inhibiting replication of SARS-CoV-2^14, 15^. However, if the exonuclease is able to remove the incorporated nucleotides including Remdesivir from this delayed termination RNA product, then the inhibitory activity of Remdesivir for SARS-CoV-2 will be reduced. Therefore, we investigated whether Pibrentasvir is efficient in preventing the excision of Remdesivir from delayed Remdesivir terminated RNA through inhibiting SARS-CoV-2 exonuclease activity. In Fig. 5, the RNA product displaying delayed Remdesivir (R) termination (sequence shown at the top of the figure) was incubated with the SARS-CoV-2 nsp14/nsp10 complex at 37 °C for 15 min in the absence (e) and presence of 20 µM of Pibrentasvir (f). MALDI-TOF MS analysis of the intact RNAs (d) and the products of the exonuclease reactions (e, f) was performed. In the absence of Pibrentasvir, exonuclease activity caused cleavage of 1-11 nucleotides from the 3’-end of the RNA as shown by the lower molecular weight fragments (e). When 20 µM Pibrentasvir was added, exonuclease activity was inhibited as shown by the reduced intensities of the fragmentation peaks and increased peak of the intact RNAs (f, 9807 Da). In Fig. 5 (a, b, c), a comparison experiment was performed with natural RNA, which produced essentially identical results. Therefore, these results indicated that the SARS-CoV-2 exonuclease activity is also significantly inhibited by Pibrentasvir to prevent nucleotide excision from RNA with delayed termination by Remdesivir.

**Fig. 5.**
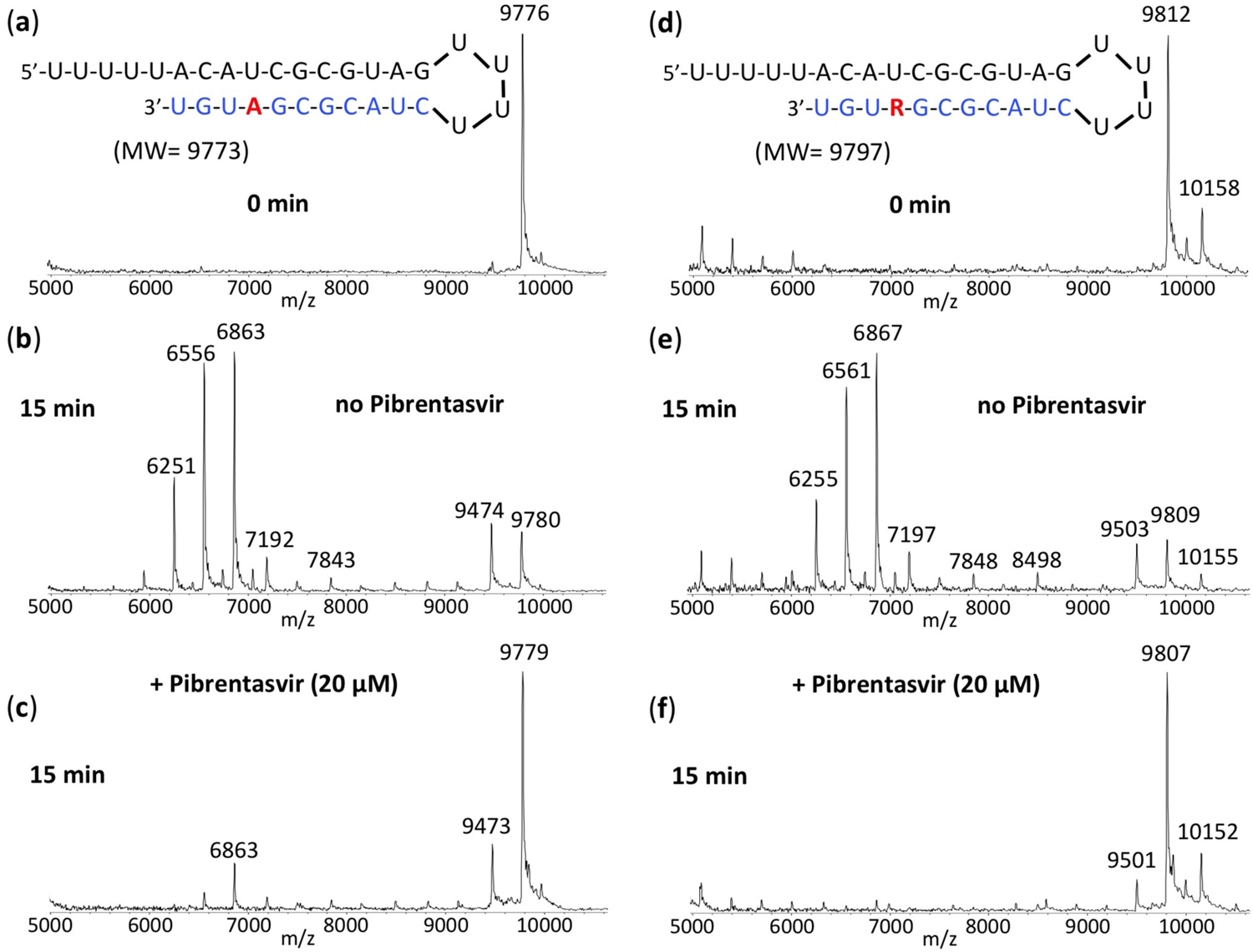
Inhibition of SARS-CoV-2 exonuclease activity by Pibrentasvir for natural RNA and RNA with delayed termination by Remdesivir (R). A mixture of 400 nM RNAs (sequences shown at the top of the figure) and 50 nM SARS-CoV-2 pre-assembled exonuclease complex (nsp14/nsp10) were incubated in buffer solution at 37 °C for 15 min in the absence (b, e) and presence of 20 µM Pibrentasvir (c, f). The intact RNAs (a, d) and the products of the exonuclease reactions (b, c, e, f) were analyzed by MALDI-TOF MS. The signal intensity was normalized to the highest peak. The peak at 9776 Da corresponds to the intact RNA (9773 Da expected) and the peak at 9812 Da corresponds to the intact Remdesivir delayed RNA termination product (9797 Da expected). The small peak at 10158 Da corresponds to mismatched incorporation of an additional G; this is likely due to the low fidelity of SARS-CoV-2 RdRp^11^. In the absence of Pibrentasvir, exonuclease activity caused nucleotide cleavage from the 3’-end of the RNA as shown by the lower molecular weight fragments corresponding to cleavage of 1-11 nucleotides (b, e). When 20 µM Pibrentasvir was added, exonuclease activity was reduced as shown by the reduced intensities of the fragmentation peaks and increased intact RNA peaks (c, f).

We previously showed that Tenofovir diphosphate (Tfv-DP) can be incorporated into RNA as an obligate terminator by the SARS-CoV-2 RdRp complex^11^. As an acyclic nucleotide, Tfv-DP lacks a normal sugar ring configuration, and thus we reasoned that it is less likely to be recognized by SARS-CoV-2 exonuclease. We therefore investigated whether Tfv can be removed by the exonuclease from the extended RNA. As shown in Fig. 6, Tfv-DP was incorporated into an RNA loop primer to produce the Tfv-terminated RNA (d). When incubation with the SARS-CoV-2 nsp14/nsp10 complex was carried out at 37 °C for 15 min, minimal cleavage was observed for Tfv-terminated RNA (e) relative to RNA extended with a natural A nucleotide (b), where there was substantial cleavage. When 20 µM Pibrentasvir was added, exonuclease activity was almost completely eliminated as shown by almost no fragmentation peaks and the essentially intact RNA peak (f). To directly compare the exonuclease resistance activities, we incubated both the RNA terminated with Tfv-DP and the natural RNA terminated with the A nucleotide in the same reaction tube, and compared the patterns obtained by MALDI-TOF MS (Fig. S-7). Because these two RNAs have different molecular weights, it was straightforward to determine whether the resulting cleavage products were derived from the Tfv- or the A-terminated RNA. The results shown in Figs. 6 and S-7 demonstrate that Tenofovir, once incorporated into RNA, is mostly resistant to excision by the SARS-CoV-2 exonuclease.

**Fig. 6.**
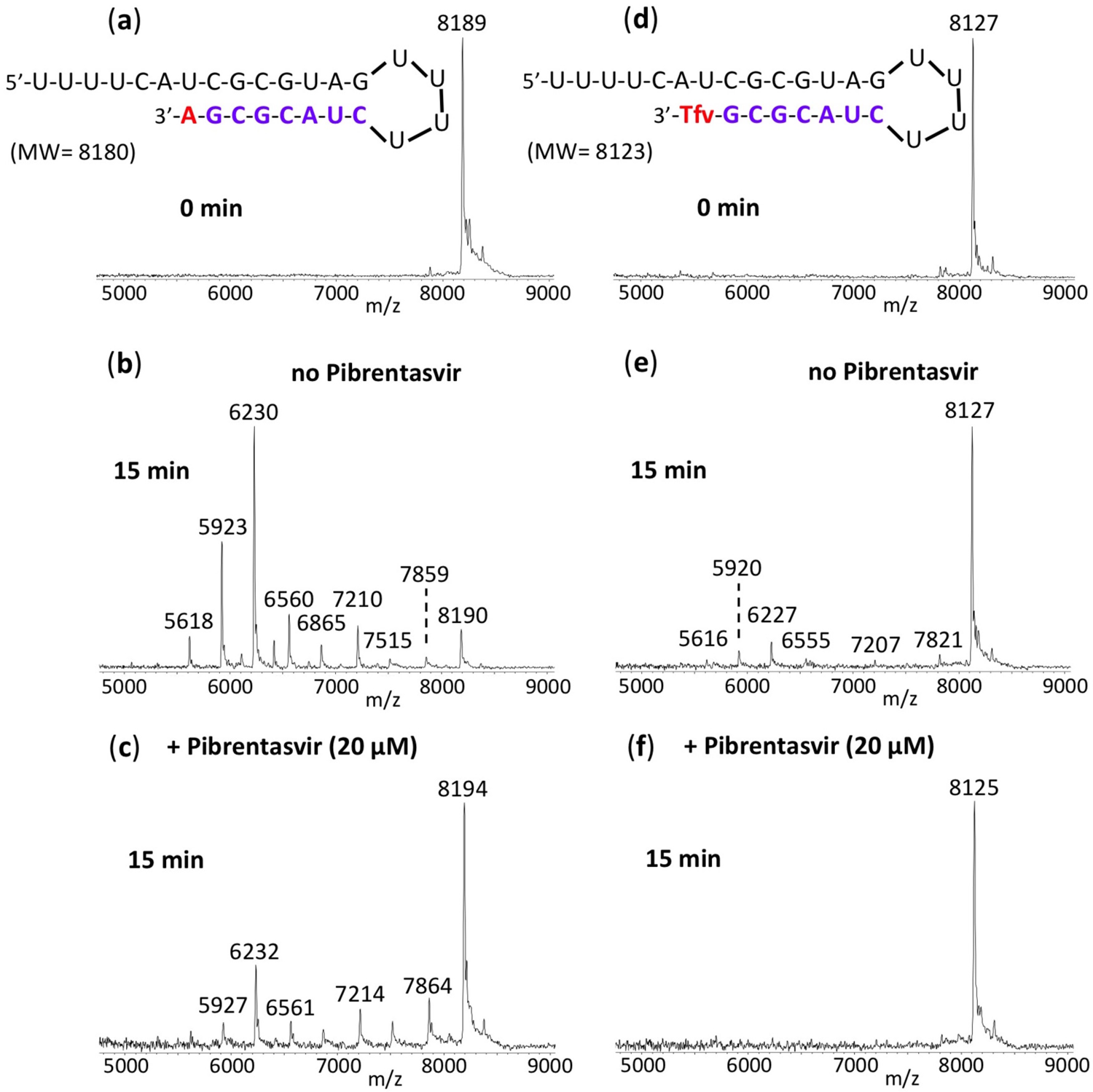
Inhibition of SARS-CoV-2 exonuclease activity by Pibrentasvir for natural RNA and Tenofovir (Tfv) terminated RNA. A mixture of 500 nM RNAs (sequences shown at the top of the figure) and 50 nM SARS-CoV-2 pre-assembled exonuclease complex (nsp14/nsp10) were incubated in buffer solution at 37 °C for 15 min in the absence (b, e) and presence of 20 µM Pibrentasvir (c, f). The intact RNAs (a, d) and the products of the exonuclease reactions (b-f) were analyzed by MALDI-TOF MS. The signal intensity was normalized to the highest peak. In the absence of Pibrentasvir, exonuclease activity caused nucleotide cleavage from the 3’-end of the natural RNA as shown by the lower molecular weight fragments corresponding to cleavage of 1-8 nucleotides (b). However, for Tfv terminated RNA, only minor cleavage was observed (e). When 20 µM Pibrentasvir was added, exonuclease activity was reduced as shown by the reduced intensities of the fragmentation peaks and increased peak height of the intact RNA (c, f).

### SARS-CoV-2 susceptibility to inhibition by immediate/delayed RNA chain terminators and error-prone nucleotide analogues is enhanced in Calu-3 cells by HCV NS5A inhibitors that inhibit the exonuclease

The prodrugs of RdRp inhibitors are activated by cellular enzymes to the active triphosphates. Therefore, the above enzymatic results need to be further evaluated at the cellular level, as interference in any of the activation pathways may impact the synergistic inhibitory activity described in the enzymatic assays. We evaluated if SARS-CoV-2’s *in vitro* susceptibility to clinically approved RdRp prodrugs could be enhanced by combining them with HCV NS5A inhibitors, which were shown to inhibit the exonuclease activity above. Calu-3 cells, which recapitulate the type II pneumocytes, cells destroyed in the course of severe COVID-19^26, 27^, were infected at a MOI of 0.1 and treated with the drugs under investigation alone or in combination.

After 2-3 days, culture supernatant was harvested and titered in VeroE6 cells to quantify the infectious viruses as plaque forming units (PFU). The NS5A inhibitors Pibrentasvir, Ombitasvir and Daclatasvir inhibited SARS-CoV-2 replication in a dose-dependent manner (Fig. S-8), but with lower potencies than Remdesivir (RDV) (Table S-1). Nevertheless, anti-SARS-CoV-2 potencies of the HCV NS5A inhibitors were 10 times higher compared to other repurposed RdRp inhibitors, such as Sofosbuvir, Tenofovir and Favipiravir (Table S-1).

To produce clinical benefit, inhibition of SARS-CoV-2 exonuclease by NS5A inhibitors need to enhance anti-coronavirus activity of RdRp inhibitors by orders of magnitude. Thus, Sofosbuvir and Tenofovir (immediate terminators), RDV (a delayed chain terminator) and Favipiravir (an error-prone nucleotide analogue) were tested at various concentrations along with the EC_25_ of Pibrentasvir (0.1 µM), Ombitasvir (0.1 µM) and Daclatasvir (0.5 µM) (Table 1 and Fig. 7). Pibrentasvir improved RDV’s efficacy at the 90% and 99% levels 6- and 3-fold, respectively (Table 1). Moreover, Pibrentasvir increased the potency of Favipiravir and Tenofovir 10-fold (Table 1). Importantly Favipiravir’s EC_90_ value becomes lower than the threshold of plasma exposure^28^ in the presence of Pibrentasvir (Table 1). Ombitasvir enhanced RDV’s and Tenofovir’s activity similarly to Pibrentasvir (Table 1). For Favipiravir, Ombitasvir resulted in a 2-log_10_ inhibition of SARS-CoV-2, reducing virus replication by 99% (Table 1 and Fig. 7D). Daclatasvir enhanced Sofosbuvir and Tenofovir potencies 10-fold at EC_50_, but only modestly increased RDV’s efficacy at EC_90_ (Table 1). In the presence of Daclatasvir, Favipiravir’s virus inhibitory efficacy at the 90% level was achievable (Table 1). These results demonstrate that the anti-exonuclease effect of NS5A inhibitors consistently enhanced the potency of prodrugs of the nucleotide analogues in the cell culture system.

**Fig. 7.**
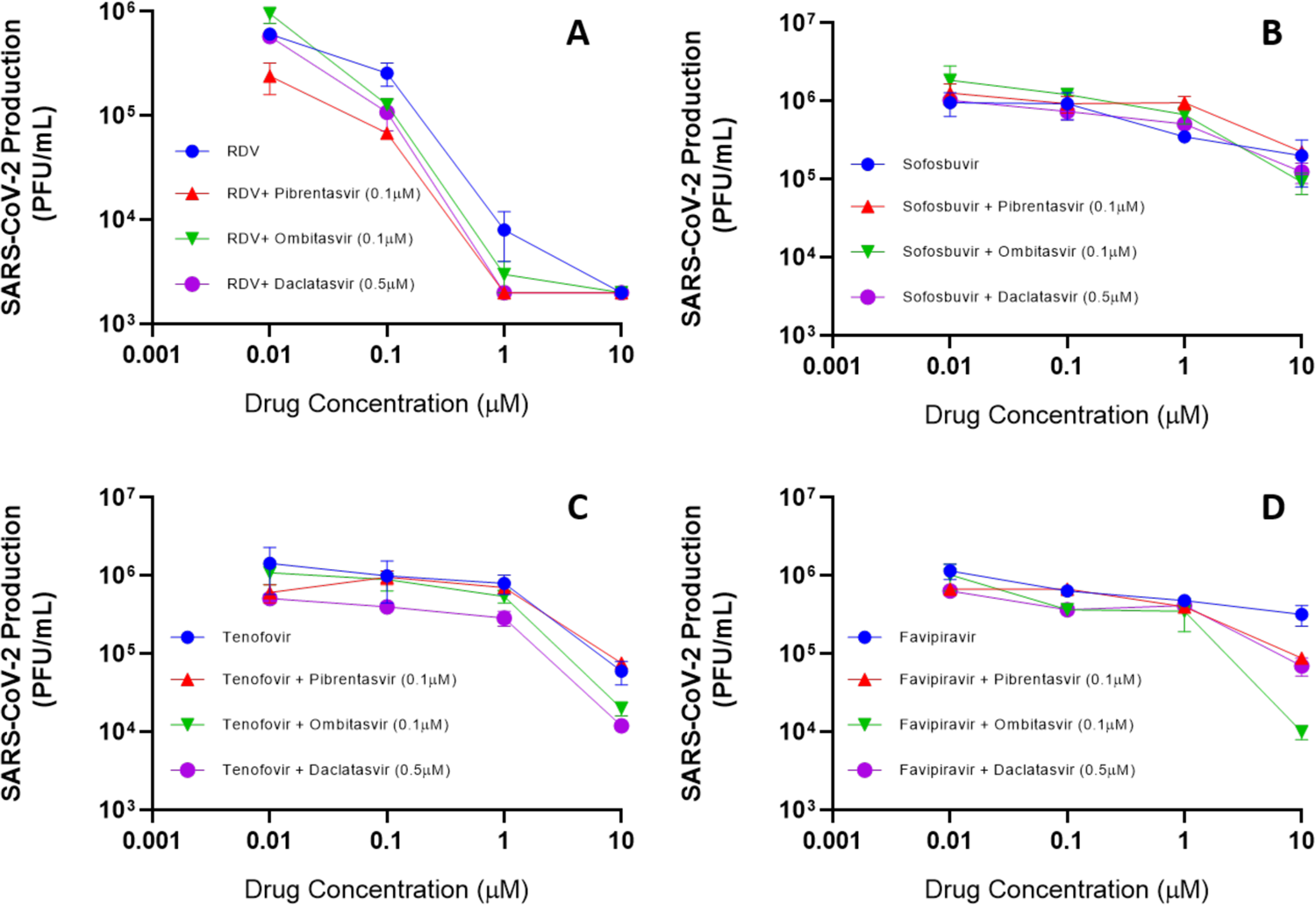
Antiviral activity of combinations of SARS-CoV-2 polymerase and exonuclease inhibitors. Calu-3 cells, at a density of 5 × 10^5^ cells/well in 48-well plates, were infected with SARS-CoV-2 at a MOI of 0.1, for 1 h at 37 °C. An inoculum was removed and cells were washed and incubated with fresh DMEM containing 2% FBS and the indicated concentration of Remdesivir (RDV) (A), Sofosbuvir (B), Tenofovir (C), and Favipiravir (D), alone and in combination with the HCV NS5A inhibitors. Supernatants were assessed after 48-72 h. Viral replication in the culture supernatant was measured as PFU/mL by titering in VeroE6 cells. Results are displayed as virus titers. The data represent means ± SEM of three independent experiments.

**Table 1.** *In vitro* pharmacological parameters of potency and efficacy of combinations of RdRp and HCV NS5A inhibitors on SARS-CoV-2 replication in Calu-3 cells

| Drug | EC<sub>50</sub> |  | EC<sub>90</sub> |  | EC<sub>99</sub> |  |
| --- | --- | --- | --- | --- | --- | --- |
|  | mean | SEM | mean | SEM | mean | SEM |
| Remdesivir (RDV) | 0.09 | 0.002 | 0.4 | 0.03 | 1.1 | 0.2 |
| Sofosbuvir | 6.2 | 0.3 | ND | ND | ND | ND |
| Tenofovir | 4.3 | 2.1 | ND | ND | ND | ND |
| Favipiravir | 7.8 | 1.2 | ND | ND | ND | ND |
| Pibrentasvir | 0.7 | 0.2 | 4.2 | 0.6 | ND | ND |
| RDV + Pibrentasvir (0.1μM) | 0.008 | 0.0009 | 0.07 | 0.03 | 0.3 | 0.09 |
| Sofosbuvir + Pibrentasvir (0.1μM) | 8.3 | 0.5 | ND | ND | ND | ND |
| Tenofovir + Pibrentasvir (0.1μM) | 0.5 | 0.05 | 8 | 1.5 | ND | ND |
| Favipiravir + Pibrentasvir (0.1μM) | 0.5 | 0.03 | 8 | 0.5 | ND | ND |
| Ombitasvir | 0.4 | 0.05 | 3.3 | 0.5 | ND | ND |
| RDV + Ombitasvir (0.1μM) | 0.008 | 0.0003 | 0.01 | 0.05 | 0.5 | 0.2 |
| Sofosbuvir + Ombitasvir (0.1μM) | 6 | 1.3 | 9 | 5 | ND | ND |
| Tenofovir + Ombitasvir (0.1μM) | 0.8 | 0.07 | 7 | 1.6 | 8.9 | 0.4 |
| Favipiravir + Ombitasvir (0.1μM) | 0.15 | 0.04 | 8 | 0.4 | 9.5 | 0.4 |
| Daclatasvir | 0.7 | 0.08 | 3.8 | 1.2 | ND | ND |
| RDV + Daclatasvir (0.5μM) | 0.008 | 0.0006 | 0.1 | 0.06 | 0.5 | 0.1 |
| Sofosbuvir + Daclatasvir (0.5μM) | 0.6 | 0.05 | 9 | 0.2 | ND | ND |
| Tenofovir + Daclatasvir (0.5μM) | 0.01 | 0.004 | 6 | 1.2 | 7.5 | 0.5 |
| Favipiravir + Daclatasvir (0.5μM) | 0.12 | 0.05 | 8 | 0.5 | ND | ND |

## Discussion

We previously demonstrated that the FDA approved HCV NS5A inhibitors, Daclatasvir and Velpatasvir, and to a lesser extent the NS5A inhibitors Elbasvir and Ledipasvir, can inhibit the SARS-CoV-2 exonuclease^29^. Of particular interest, Daclatasvir and Velpatasvir inhibit both the SARS-CoV-2 polymerase and exonuclease^20, 21^. Here, we showed that two additional NS5A inhibitors, Pibrentasvir and Ombitasvir, also inhibit the exonuclease, and have the highest inhibitory activity based on our molecular assay. These compounds are predicted to interfere with the binding of the Mg^++^ ion with the 3’ terminus of the RNA in the active site of the exonuclease (nsp14). The Mg^++^ ion coordinates amino acid residues Asp-90, Glu-92, Glu-191 and Asp-273 and the 3’ terminus of the RNA. Because the NS5A inhibitors interfere with this coordination, they are likely to prevent nucleotide excision from the RNA (Fig. 2). A recent *in silico* modeling study has suggested that Ritonavir also binds to the active site of nsp14, which led the authors to the prediction that Ritonavir may inhibit exonuclease activity^30^. We have experimentally shown (Fig. S-9) that Ritonavir and Lopinavir, HIV protease inhibitors that make up the combination drug Kaletra, inhibit the SARS-CoV-2 exonuclease in a concentration-dependent manner, but with less potency than Pibrentasvir and Ombitasvir. Kaletra has gone through extensive clinical trials for COVID-19 but shows limited efficacy by itself^31, 32^. Recently, using a digital drug development approach combining artificial intelligence and experimental validation to screen drug combinations for potential combination therapy against SARS-CoV-2, it was determined that one of the most effective potential combinations was Remdesivir, Ritonavir and Lopinavir^33^. This synergy of Ritonavir and Lopinavir with Remdesivir may be due to the exonuclease inhibitory activity of these two drugs (Fig. S-9).

NS5A inhibitors enhance the antiviral activity of RDV, Tenofovir, Sofosbuvir and Favipiravir, likely due to their inhibition of exonuclease activity. The combinations of drugs described here, especially the orally available antivirals, have the potential to progress to COVID-19 clinical trials. We show that by combining Pibrentasvir or Ombitasvir with Remdesivir, Sofosbuvir, Tenofovir or Favipiravir, higher inhibitory activity for SARS-CoV-2 was achievable at lower doses, bringing the nucleotides’ pharmacological parameters more in line with their pharmacokinetic exposures^28, 34, 35^. The nucleotide inhibitors Remdesivir and Sofosbuvir are incorporated into the replicating RNA to inhibit further polymerase reaction, but they are rapidly removed by the exonuclease in the absence of exonuclease inhibitors such as Pibrentasvir and Ombitasvir. However, in the presence of these inhibitors, any nucleotide inhibitors incorporated into the RNA will not be rapidly excised by the exonuclease and their inhibitory effect on overall viral RNA replication will be substantially enhanced. Recently, the coronavirus exonuclease has also been shown to promote viral recombination which can increase the emergence of novel strains^36^, providing additional incentive for targeting this enzyme with antivirals.

Previously, Baddock *et al.* screened a series of molecules for SARS-CoV-2 exonuclease activity, including Ebselen and Disulfiram, with Ebselen having the higher inhibitory activity^37^. Ebselen and Disulfiram have multiple additional SARS-CoV-2 protein targets and Chen *et al.* used a combination of Ebselen/Disulfiram with Remdesivir to demonstrate a small synergistic inhibitory effect on viral replication^38^. Recently, it has been reported that HCV nsp3/4A protease inhibitors that inhibit the SARS-CoV-2 papain-like protease synergize with Remdesivir to suppress viral replication in cell culture^39^. In our investigation, by focusing on combinations of clinically approved drugs used during routine treatment of HIV/HCV/influenza infected individuals, we discovered significant synergistic combinations of drugs that inhibit both the SARS-CoV-2 polymerase and exonuclease activities.

Favipiravir (6-fluoro-3-hydroxy-2-pyrazinecarboxamide, Fig. 1c), a drug used to treat influenza, has been investigated for treatment of COVID-19^40, 41^. It has a modified nucleobase that can base pair with both C and U. Favipiravir is converted to the active triphosphate form, Favipiravir-ribofuranosyl-5’-triphosphate (Favipiravir-RTP), by a cellular enzyme, hypoxanthine guanine phosphoribosyltransferase and cellular kinases (Fig. S-10). Favipiravir-RTP acts as a viral RNA polymerase inhibitor, with great potential in the treatment of a wide variety of RNA virus infections, including different strains of influenza^42, 43^. Favipiravir-RTP is incorporated into the growing RNA chain by the SARS-CoV-2 RdRp complex and causes C-to-U and G-to-A transitions. In a more recent study, Favipiravir was found to exhibit antiviral activity against SARS-CoV-2 via a combination of mechanisms, including slowing RNA synthesis, initiation of delayed chain termination, and lethal mutagenesis^23, 40^. The 3-D structure of the SARS-CoV-2 RdRp complex (nsp12, nsp7 and nsp8) with the RNA substrate and Favipiravir-RTP was determined, providing insight into the mode of action of this drug^44, 45^. Here, we demonstrated that the excision of Favipiravir from the 3’ end of RNA by the SARS-CoV-2 exonuclease complex is greatly reduced in the presence of Pibrentasvir (Fig. S-4). Since Pibrentasvir and Ombitasvir inhibited the SARS-CoV-2 exonuclease proofreading activity, when these drugs are combined with Favipiravir, it is more likely that Favipiravir-induced mutations will persist and jeopardize viral replication. This is supported by our cell culture virus inhibition data indicating that the combination of Favipiravir with Ombitasvir resulted in a 2-log_10_ inhibition, reducing virus replication by 99% (Table 1).

N^4^-hydroxycytidine (NHC, Fig. 1e) has been shown to inhibit SARS-CoV-2 and other coronaviruses in mice and human airway epithelial cells^24^. NHC is incorporated opposite either A or G in the template strand. Molnupiravir, which is activated to the triphosphate form (NHC-TP) by cellular enzymes, is currently in a COVID-19 clinical trial^46^. Here, we showed that the excision of NHC from the 3’ end of RNA by the SARS-CoV-2 exonuclease complex is substantially reduced in the presence of Pibrentasvir (Fig. S-5).

AT-527 (Fig. 1f), a prodrug of a guanosine nucleotide analogue with both the base and phosphate group masked, is converted by cellular enzymes to the active triphosphate form, 2’-fluoro-2’-methyl guanosine-5’-triphosphate (AT-9010, G_fm_-triphosphate)^25^. The active triphosphate form of AT-527 is structurally similar to that of Sofosbuvir with the only difference being that it has a guanine base in place of uracil. AT-527 inhibits SARS-CoV-2 by targeting both the RdRp activity and the Nidovirus RdRp-associated nucleotidyltransferase (NiRAN) activity of nsp12, which are essential for viral RNA replication and transcription^47^. AT-527 is currently in COVID-19 clinical trials. Here, we demonstrated that the excision of AT-527 from the 3’ end of RNA by the SARS-CoV-2 exonuclease is largely abrogated by the exonuclease inhibitor Pibrentasvir (Fig. S-6).

Additionally, we have shown that when Tenofovir is incorporated into RNA, it is mostly resistant to excision by the SARS-CoV-2 exonuclease, which might be explained by the fact that it is an acyclic nucleotide analogue, interfering with its recognition by the exonuclease. The addition of Pibrentasvir almost completely protected the Tenofovir terminated RNA from excision by exonuclease. The combination of NS5A inhibitors with Tenofovir also substantially increased the virus inhibition efficiency in cell culture (Table 1).

Our results from the molecular enzymatic assays and the virus cell culture inhibitory data both indicate that the use of inhibitors of the viral RdRp and the exonuclease proofreader as a combination therapy is expected to have enhanced efficacy in treating COVID-19. This combination approach may allow two-log_10_ reduction of virus replication using concentrations consistent with approved plasma exposure. We anticipate that the various nucleotide analogue inhibitors of the SARS-CoV-2 RdRp that are in clinical trials for COVID-19 will benefit from the addition of exonuclease inhibitors.

## Methods

The RdRp of SARS-CoV-2, referred to as nsp12, and its two protein cofactors, nsp7 and nsp8, shown to be required for the processive polymerase activity of nsp12, were cloned and purified as described^11, 20^. The 3′-exonuclease, referred to as nsp14, and its protein cofactor, nsp10, were cloned and expressed based on the SARS-CoV-2 genome sequence. Remdesivir triphosphate (RDV-TP) and NHC triphosphate were purchased from MedChemExpress (Monmouth Junction, NJ), Sofosbuvir triphosphate (SOF-TP) was purchased from Sierra Bioresearch (Tucson, AZ), Tenofovir diphosphate (Tfv-DP) was purchased from Alfa Chemistry (Ronkonkoma, NY), and Favipiravir triphosphate and AT-9010 were purchased from NuBlocks LLC (Oceanside, CA). Pibrentasvir was purchased from Cayman Chemical Company (Ann Arbor, MI) and Ombitasvir, Daclatasvir, Ritonavir and Lopinavir were purchased from Selleckchem (Houston, TX). UTP, GTP and ATP were purchased from Fisher Scientific. The RNA oligonucleotides (template-loop-primers) were purchased from Dharmacon (Horizon Discovery, Lafayette, CO).

### Expression and purification of the SARS-CoV-2 exonuclease nsp14/nsp10 complex

The 3′-exonuclease, referred to as nsp14, and its protein cofactor, nsp10, were cloned and expressed based on the SARS-CoV-2 genome sequence. The pRSFDuet-1 plasmids (Novagen) encoding SARS-CoV-2 nsp14 or nsp10 engineered with an N-terminal His-SUMO tag were prepared as follows: SARS-CoV-2 RNA isolated from the supernatant of SARS-CoV-2-infected Vero E6 cells was provided by Benjamin R. tenOever^48^. The portion of the RNA encoding nsp10 and nsp14 was reverse transcribed into cDNA using gene-specific primers (see below) and SuperScript III Reverse Transcriptase (Thermo Fisher). The nsp10 and nsp14 coding sequences were PCR amplified using forward and reverse gene specific primers flanked by *Bam*HI and *Xho*I recombination sites, respectively. PCR products were digested with *Bam*HI and *Xho*I and subsequently ligated into *Bam*HI and *Xho*I-digested pRSFDuet-His6-sumo vector and sequence verified. pRSFDuet-His6-sumo is a modified pRSFDuet-1 vector (Novagen) bearing an N-terminal His6-SUMO-tag cleavable by the ubiquitin-like protease (ULP1).

List of PCR primers used:

NSP14_pRSF_BamHI_fw CGCGGATCCGCTGAAAATGTAACAGGACTCTTTAAA

NSP14_pRSF_XhoI_rev CCCGCTCGAGCGGTCACTGAAGTCTTGTAAAAGTGTTCCAGAGG

RT_primer_NSP14 TTCTTGGCTATGTCAGTCATAGAACAAAC

NSP10_pRSF_BamHI_fw CGCGGATCCGCTGGTAATGCAACAGAAGTGCCTGCC

NSP10_pRSF_XhoI_rev CCCGCTCGAGCGGTCACTGAAGCATGGGTTCGCGGAGTTGATC

RT_primer_NSP10 GATGTTGATATGACATGGTCGTAACAGC

The nsp14 and nsp10 proteins were expressed in *Escherichia coli* BL21-CodonPlus(DE3)-RIL (Stratagene). The bacteria were grown in Luria–Bertani medium supplemented with 50 mg/mL kanamycin at 37 °C to an OD600 of 0.6, and induced with 0.4 mM isopropyl β-D-1-thiogalactopyranoside and 50 µM ZnCl_2_ overnight at 18 °C. Cells were collected via centrifugation at 5,000 × g and equal volumes of the nsp14 and nsp10 bacterial cells were then mixed for nsp14-nsp10 protein complex purification. The cells were lysed via sonication in Lysis Buffer (500 mM NaCl, 20 mM imidazole, 20 mM Tris-HCl, pH 8.0, 1 mM phenylmethylsulfonyl fluoride). After centrifugation at 40,000 × g, the supernatant was loaded onto 5 mL Nickel Sepharose 6 fast flow resins (GE Healthcare) in a gravity flow column. The target protein was eluted using Lysis Buffer supplemented with 500 mM imidazole. The eluted protein was incubated with ULP1 (lab stock) during dialysis at 4 °C overnight against a buffer containing 20 mM Tris-HCl, pH 7.5, 20 mM imidazole, 150 mM NaCl, 100 µM ZnCl_2_, and 5mM β-mercaptoethanol. Then the sample was loaded onto the HisTrap FF column (GE Healthcare) to remove the His-SUMO tag, and the flow-through was collected. The target proteins were further purified through a Superdex200 10/300 gel filtration column (GE Healthcare) in a buffer containing 20 mM HEPES, pH 7.4, 150 mM NaCl, 1 mM MgCl_2_, and 1 mM dithiothreitol. The fractions corresponding to the nsp14/nsp10 complex were detected by SDS-PAGE and collected. The protein sample was flash-frozen in liquid nitrogen and stored at −80 °C.

### Extension reactions with SARS-CoV-2 RNA-dependent RNA polymerase complex to produce Remdesivir (RDV) and Sofosbuvir (SOF) terminated RNAs

10 µL of 10 µM RNA template-loop-primers (5’-UUUUCAUCGCGUAGUUUUCUACGCG-3’ for RDV-TP extension; 5’-UUUUCUACGCGUAGUUUUCUACGCG-3’ for SOF-TP extension) in 1 × RdRp reaction buffer was annealed by heating to 75 °C for 3 min and cooling to room temperature. 5 µL of 8 µM RdRp complex (nsp12/nsp7/nsp8)^11, 20^ in 1 × reaction buffer was added to the annealed RNA template-loop-primer solution and incubated for an additional 10 min at room temperature. Finally, 5 µL of a solution containing 0.2 mM RDV-TP or 2 mM SOF-TP in 1 × reaction buffer was added and incubation was carried out for 2 h at 30 °C. The final concentrations of reagents in the 20 µL extension reactions were 2 µM nsp12/nsp7/nsp8, 5 µM RNA template-loop-primer, 50 µM RDV-TP and 500 µM SOF-TP. The 1 × reaction buffer contains the following reagents: 10 mM Tris-HCl pH 8, 10 mM KCl, 2 mM MgCl_2_ and 1 mM β-mercaptoethanol. Desalting of the reaction mixture was performed with an Oligo Clean & Concentrator kit (Zymo Research) resulting in ∼10 µL purified aqueous RNA solutions. 1 µL of each solution was subjected to MALDI-TOF MS (Bruker ultrafleXtreme) analysis. The remaining ∼9 µL extended template-loop-primer solutions were used to test exonuclease activity.

### Extension reactions with Therminator II to produce AT-9010 terminated RNA

The RNA template-loop primer (5’-UUUUCUCCGCGUAGUUUUCUACGCG-3’) was annealed at 75 °C for 3 minutes and cooled to room temperature for 25 minutes in 1 × ThermoPol buffer before adding the other ingredients. The final 20 µL mixture containing 5 µM of the RNA template-loop primer, 250 µM, 500 µM or 1 mM AT-9010, 2 mM MnCl_2_ and 0.2 unit Therminator II in 1 × ThermoPol buffer was incubated in a thermal cycler using the following protocol (28 cycles of 45 °C for 30 sec, 55 °C for 30 sec, 65°C for 30 sec). Desalting of the reaction mixture was performed with an Oligo Clean & Concentrator kit (Zymo Research) resulting in ∼10 µL purified aqueous RNA solutions. 1 µL of each solution was subjected to MALDI-TOF MS (Bruker ultrafleXtreme) analysis. The remaining ∼9 µL extended template-loop-primer solutions were used to test exonuclease activity.

### Extension reactions with SARS-CoV-2 RNA-dependent RNA polymerase to produce Favipiravir or NHC terminated RNAs

10 µL of 10 µM RNA template-loop-primers (5’-UUUUCAUCGCGUAGUUUUCUACGCG-3’ for Fav-RTP extension opposite U; 5’-UUUUCACCGCGUAGUUUUCUACGCG-3’ for Fav-RTP extension opposite C; 5’-UUUUCUACGCGUAGUUUUCUACGCG-3’ for NHC-TP extension opposite A; 5’-UUUUCUGCGCGUAGUUUUCUACGCG-3’ for NHC-TP extension opposite G) in 1 × RdRp reaction buffer was annealed by heating to 75 °C for 3 min and cooling to room temperature. 5 µL of 12 µM RdRp complex (nsp12/nsp7/nsp8)^11, 20^ in 1 × reaction buffer was added to the annealed RNA template-loop-primer solution and incubated for an additional 10 min at room temperature. Finally, 5 µL of a solution containing ∼2 mM Fav-RTP or 0.4 mM NHC in 1 × reaction buffer was added and incubation was carried out for 3 h at 30 °C. The final concentrations of reagents in the 20 µL extension reactions were 3 µM nsp12/nsp7/nsp8, 5 µM RNA template-loop-primer and ∼500 µM Fav-RTP or 100 µM NHC-TP. The 1 × reaction buffer contains the following reagents: 10 mM Tris-HCl pH 8, 10 mM KCl, 2 mM MgCl_2_ and 1 mM β-mercaptoethanol. Desalting of the reaction mixture was performed with an Oligo Clean & Concentrator kit (Zymo Research) resulting in ∼10 µL purified aqueous RNA solutions. 1 µL of each solution was subjected to MALDI-TOF MS (Bruker ultrafleXtreme) analysis. The remaining ∼9 µL extended template-loop-primer solutions were used to test exonuclease activity.

### Extension reactions with reverse transcriptase to produce Tenofovir (Tfv) terminated RNAs

The RNA template-loop primer (5’-UUUUCAUCGCGUAGUUUUCUACGCG-3’) was annealed at 75 °C for 3 minutes and cooled to room temperature for 25 minutes before adding the remaining ingredients. The final 20 µL mixture containing 5 µM of the RNA template-loop primer, 500 µM Tenofovir-DP and 200 units Superscript IV Reverse Transcriptase in 1 × reaction buffer (50 mM Tris-HCl (pH 8.3), 4 mM MgCl_2_, 10 mM DTT, 50 mM KCl) was incubated at 45 °C for 3 hr. Desalting of the reaction mixture was performed with an Oligo Clean & Concentrator kit (Zymo Research) resulting in ∼10 µL purified aqueous RNA solutions. 1 µL of each solution was subjected to MALDI-TOF MS (Bruker ultrafleXtreme) analysis. The remaining ∼9 µL extended template-loop-primer solutions were used to test exonuclease activity.

### Extension reactions with SARS-CoV-2 RNA-dependent RNA polymerase to produce Remdesivir (RDV) delayed terminated RNA

10 µL of 10 µM RNA template-loop-primers (5’-UUUUUACAUCGCGUAGUUUUCUACGCG-3’ for RDV-TP + UTP extension) in 1 × RdRp reaction buffer was first annealed by heating to 75 °C for 3 min and cooling to room temperature. 5 µL of 8 µM RdRp complex (nsp12/nsp7/nsp8)^11, 20^ in 1 × reaction buffer was added to the annealed RNA template-loop-primer solution and incubated for an additional 10 min at room temperature. 5 µL of a solution containing 0.2 mM RDV-TP and 0.2 mM UTP in 1 × reaction buffer was added and incubation was carried out for 1 h at 30 °C. The final concentrations of reagents in the 20 µL extension reactions were 2 µM nsp12/nsp7/nsp8, 5 µM RNA template-loop-primer, 50 µM RDV-TP and 50 µM UTP. The composition of 1 × reaction buffer was as described above. Desalting of the reaction mixture was performed with an Oligo Clean & Concentrator kit (Zymo Research) resulting in ∼10 µL purified aqueous RNA solutions. 1 µL of each solution was subjected to MALDI-TOF MS (Bruker ultrafleXtreme) analysis. The remaining ∼9 µL extended template-loop-primer solutions were used for the second extension reaction.

Then, 10 µL of ∼16 µM Remdesivir (R) and uridine extended RNA template-loop-primers (5’-UUUUUACAUCGCGUAGUUUUCUACGCGRU-3’ for second extension) in 1 × RdRp reaction buffer was annealed by heating to 75 °C for 3 min and cooling to room temperature. 5 µL of 8 µM RdRp complex (nsp12/nsp7/nsp8)^11, 20^ in 1 × reaction buffer was added to the annealed RNA template-loop-primer solution and incubated for an additional 10 min at room temperature. 5 µL of a solution containing 80 µM GTP and 80 µM UTP in 1 × reaction buffer was added and incubation was carried out for 1 h at 30 °C. The final concentrations of reagents in the 20 µL extension reactions were 2 µM nsp12/nsp7/nsp8, ∼4 µM RNA template-loop-primer, 20 µM GTP and 20 µM UTP. The composition of 1 × reaction buffer was as described above. Desalting of the reaction mixture was performed with an Oligo Clean & Concentrator kit (Zymo Research) resulting in ∼10 µL purified aqueous RNA solutions. 1 µL of each solution was subjected to MALDI-TOF MS (Bruker ultrafleXtreme) analysis. The remaining ∼9 µL extended template-loop-primer solutions were used to test exonuclease activity.

### SARS-CoV-2 exonuclease reactions in the presence and absence of Pibrentasvir, Ombitasvir, Daclatasvir, Ritonavir and Lopinavir

The U-terminated RNA, A-terminated RNA, G-terminated RNA, the Remdesivir (RDV), Sofosbuvir (SOF), Favipiravir (Fav), N^4^-hydroxycytidine (Nhc), AT-9010 and Tenofovir (Tfv) terminated RNAs as well as the RDV delayed terminated RNA products from above, were annealed by heating to 75 °C for 3 min and cooling to room temperature in 1 × exonuclease reaction buffer. To a 14 µL solution of 71.4 nM exonuclease complex (nsp14/nsp10) in 1 × exonuclease reaction buffer, 1 µL of DMSO with or without various concentrations of Pibrentasvir was added and incubated for 15 min at room temperature. Then, 5 µL of the annealed RNA (2 µM) in 1 × exonuclease reaction buffer was added to the exonuclease/Pibrentasvir mixture and incubated at 37 °C for 15 min. The final concentrations of reagents in the 20 µL reactions were 50 nM nsp14/nsp10, 500 nM RNA, 0-20 µM Pibrentasvir and 5% DMSO. The 1 × exonuclease reaction buffer contains the following reagents: 40 mM Tris-HCl pH 8, 1.5 mM MgCl_2_ and 5 mM DTT. After incubation for 15 min, each reaction was quenched by adding 2.2 µL of an aqueous solution of EDTA (100 mM). Following desalting using an Oligo Clean & Concentrator (Zymo Research), the samples were subjected to MALDI-TOF MS (Bruker ultrafleXtreme) analysis. SARS-CoV-2 exonuclease reactions in the presence of Ombitasvir, Daclatasvir, Ritonavir and Lopinavir were conducted analogously to the experiments with Pibrentasvir.

### Comparison of SARS-CoV-2 exonuclease reaction for Tenofovir terminated RNA with natural RNA

The A-terminated RNA (2 µM) and Tenofovir extended RNA (2 µM) (sequences shown in Fig. S-4) were annealed by heating to 75 °C for 3 min and cooling to room temperature in 1 × exonuclease reaction buffer. After annealing, the two RNAs were mixed in equal volumes. To a 10 µL solution of 100 nM exonuclease complex (nsp14/nsp10) in 1 × exonuclease reaction buffer 10 µL annealed RNA mixture (1 µM each) was added and incubated at 37 °C for 15 min. The final concentrations of reagents in the 20 µL reactions were 50 nM nsp14/nsp10, ∼500 nM A-terminated RNA and ∼500 nM Tenofovir terminated RNA. After incubation for 15 min, each reaction was quenched by adding 2.2 µL of an aqueous solution of EDTA (100 mM). Following desalting using an Oligo Clean & Concentrator (Zymo Research), the samples were subjected to MALDI-TOF MS (Bruker ultrafleXtreme) analysis.

### Molecular docking procedure

The chemical structures for Pibrentasvir, Ombitasvir, Elbasvir, Daclatasvir, Ledipasvir and Velpatasvir (Fig. S-1) were obtained from PubChem^49^ and minimized in terms of energy by Density Functional Theory (DFT), with the Becke-3-Lee Yang Parr (B3LYP) method and the standard 6-31G* basis set, available in Spartan’18 software (Wavefunction, Inc., Irvine, USA)^50^. Since the 3D structure for the SARS-CoV-2 nsp-14 exonuclease was not available, a structural model was built via the on-line tool Swiss Model software (University of Basel, Basel, Switzerland)^51^ using the crystallographic structure of SARS-CoV nsp-14 as template (Protein Data Bank (PDB) code: 5C8T)^52^ and the following SARS-CoV-2 nsp-14 amino acid sequence: AENVTGLFKDCSKVITGLHPTQAPTHLSVDTKFKTEGLCVDIPGIPKDMTYRRLISMMGFKMNYQVNGYPNMFITREEAIRHVRAWIGFDVEGCHATREAVGTNLPLQLGFSTGVNLVAVPTGYVDTPNNTDFSRVSAKPPPGDQFKHLIPLMYKGLPWNVVRIKIVQMLSDTLKNLSDRVVFVLWAHGFELTSMKYFVKIGPERTCCLCDRRATCFSTASDTYACWHHSIGFDYVYNPFMIDVQQWGFTGNLQSNHDLYCQVHGNAHVASCDAIMTRCLAVHECFVKRVDWTIEYPIIGDELKINAACRKVQHMVVKAALLADKFPVLHDIGNPKAIKCVPQADVEWKFYDAQPCSDKAYKIEELFYSYATHSDKFTDGVCLFWNCNVDRYPANSIVCRFDTRVLSNLNLPGCDGGSLYVNKHAFHTPAFDKSAFVNLKQLPFFYYSDSPCESHGKQVVSDIDYVPLKSATCITRCNLGGAVCRHHANEYRLYLDAYNMMISAGFSLWVYKQFDTYNLWNTFTRLQ^53^. About five models were generated based on the standard ‘auto model’ routine of Swiss Model software. The best modeled structure was chosen according to the Qualitative Model Energy Analysis (QMEAN) and Global Model Quality Estimation (GMQE) values.

The molecular docking calculations were performed using GOLD 2020.2 software (Cambridge Crystallographic Data Centre, Cambridge, UK)^54^. Hydrogen atoms were added to the protein structure according to the data inferred by software on the ionization and tautomeric states. Since there are no crystallographic structures for drugs associated with SARS-CoV-2 nsp14 exonuclease, the standard function *ChemPLP* was used for the molecular docking calculations, selecting an 8 and 10 Å radius spherical cavity around the active binding site. The figures for the docking poses of the largest docking score value was generated with PyMOL Delano Scientific LLC software (Schrödinger, New York, USA)^55^.

### Cells and Virus

African green monkey kidney cells (Vero, subtype E6) and the human lung epithelial cell line (Calu-3 cells) were cultured in high glucose DMEM and low glucose DMEM medium, both complemented with 10% FBS, 100 U/mL penicillin and 100 µg/mL streptomycin at 37 °C in a humidified atmosphere with 5% CO_2_. SARS-CoV-2 was prepared in Vero E6 cells at a MOI of 0.01. Originally, the isolate was obtained from a nasopharyngeal swab from a confirmed case in Rio de Janeiro, Brazil (GenBank #MT710714; Institutional Review Broad approval, 30650420.4.1001.0008). All procedures related to virus culture were handled in a Biosafety Level 3 (BSL3) multiuser facility according to WHO guidelines. Virus titers were determined as plaque forming units (PFU)/mL. Virus stocks were kept in −80°C freezers.

### Cytotoxicity assay

Monolayers of 1.5 x 10^4^ cells in 96-well plates were treated for 3 days with various concentrations (semi-log dilutions from 1000 to 10 µM) of the antiviral drugs. Then, 5 mg/ml 2,3-bis-(2-methoxy-4-nitro-5-sulfophenyl)-2*H*-tetrazolium-5-carboxanilide (XTT) in DMEM was added to the cells in the presence of 0.01% *N*-methyl dibenzopyrazine methyl sulfate (PMS). After incubating for 4 h at 37 °C, the cells were measured in a spectrophotometer at 492 nm and 620 nm. The 50% cytotoxic concentration (CC_50_) was calculated by a non-linear regression analysis of the dose response curves.

### Yield-reduction assay

Calu-3 cells (5 x 10^5^ cells/well) in 48-well plates were infected at a MOI of 0.1 for 1 h at 37 °C. The cells were washed, and various concentrations of compounds were added to DMEM with 10% FBS. 48-72 h supernatants were collected and the harvested virus was quantified as PFU/mL by titering in Vero E6 cells. A variable slope non-linear regression analysis of the dose response curves was performed to calculate the concentration at which each drug inhibited the virus production by 50% (EC_50_), 90% (EC_90_) and 99% (EC_99_).

### Virus titration

Monolayers of Vero E6 cells (2 x 10^4^ cells/well) in 96-well plates were infected with serial dilutions of supernatants containing SARS-CoV-2 for 1 h at 37 °C. Fresh semi-solid medium containing 2.4% CMC was added and the culture was maintained for 72 h at 37 °C. Cells were fixed with 10% formalin for 2 h at room temperature and then stained with crystal violet (0.4%). Plaque numbers were scored in at least 3 replicates per dilution by independent readers, who were blinded with respect to the source of the supernatant. The virus titers were determined as PFU/mL.

## Statistical analysis

The assays were performed blinded by one professional, codified and then read by another professional. All experiments were carried out at least three independent times, including a minimum of two technical replicates in each assay. The dose-response curves used to calculate pharmacological parameters were generated by variable slope plot from Prism GraphPad software 8.0. The equations to fit the best curve were generated based on R^2^ values ≥ 0.9.

## Acknowledgements

This work was supported by the Jack Ma Foundation, a gift from Columbia Engineering Member of the Board of Visitors Dr. Bing Zhao, and Fast Grants (to Jingyue Ju), the Maloris Foundation and the Memorial Sloan-Kettering Core Grant (P30CA008748) (to Dinshaw J. Patel), a grant from The JPB Foundation to Rockefeller University (to Thomas Tuschl). Funding was also provided by Conselho Nacional de Desenvolvimento Científico e Tecnológico (CNPq), Fundação de Amparo à Pesquisa do Estado do Rio de Janeiro (FAPERJ) and Coordenação de Aperfeiçoamento de Pessoal de Nível Superior - Brasil (CAPES) - Finance Code 001 (to Thiago Moreno L. Souza and Patricia T. Bozza). CNPq, CAPES and FAPERJ also support the National Institutes of Science and Technology Program (INCT-IDPN). Oswaldo Cruz Foundation/FIOCRUZ supports this study under the auspices of the Inova Program (B3-Bovespa funding) (to Thiago Moreno L. Souza). Dr. Andre Sampaio from Farmanguinhos, platform RPT11M, is acknowledged for kindly donating the Calu-3 cells. We thank Dr. Andrew Owen from the University of Liverpool for insightful discussions.

## Author contributions

X.W., C.Q.S., S.J., O.A.C, C.T., N. F.-R., M.C., J.R.T., W.X., C.M. and A.G. performed the experiments. X.W., C.Q.S., S.J., C.T., N.F.-R., M.C., J.R.T., P.T.B., X.L., S.K., J.J.R., T.M.L.S. and J.J. analyzed the data. X.W., C.Q.S., S.J., C.T., M.C., J.R.T., X.L., S.K., P.T.B., J.J.R., T.M.L.S. and J.J. prepared the manuscript. C.Q.S., D.J.P., T.T., T.M.L.S. and J.J. conceptualized the experiments.

## Additional information

Correspondence and requests for materials should be addressed to J.J. and T.M.L.S.

## SUPPLEMENTARY INFORMATION

**Fig. S-1.**
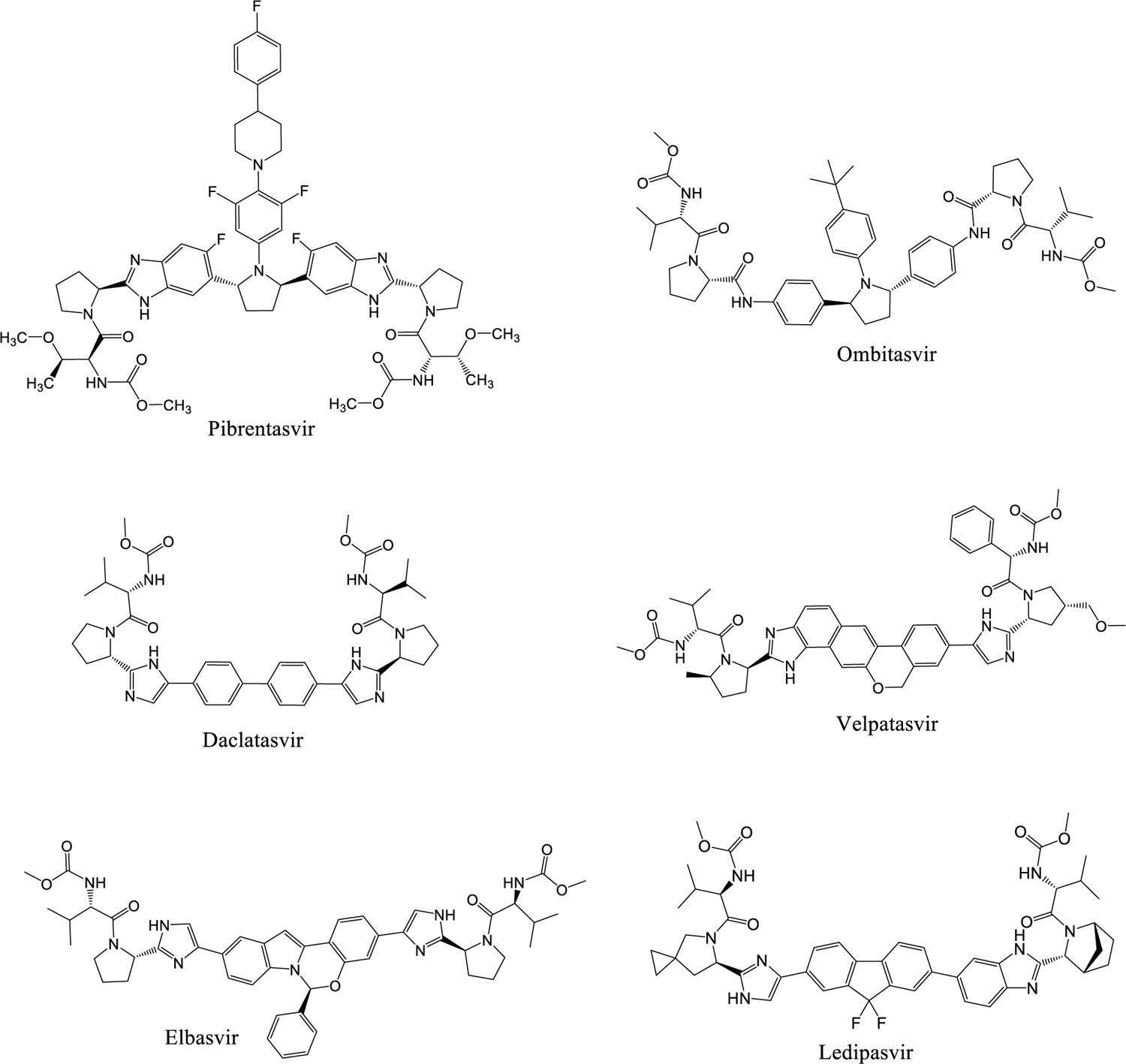
Structures of HCV NS5A inhibitors: Pibrentasvir, Ombitasvir, Daclatasvir, Velpatasvir, Elbasvir and Ledipasvir.

**Fig. S-2.**
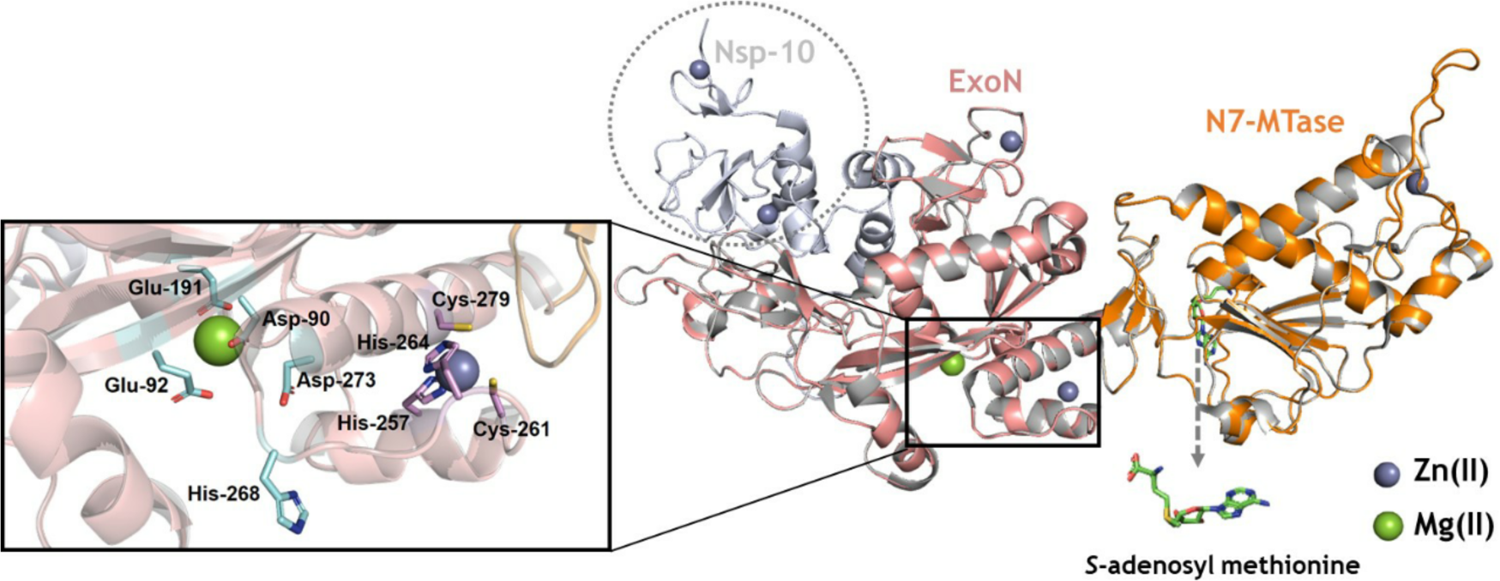
Superposition of the SARS-CoV nsp14 (PDB code: 5C8T, in gray) and SARS-CoV-2 nsp14 model (beige and orange for ExoN and N7-MTase domains, respectively). Selected amino acid residues in the exonuclease catalytic site are represented as stick representations in cyan. The co-substrate involved in methyltransferase, *S*-adenosyl methionine (SAM), is represented using a stick model in green. The hydrogen atoms were omitted for better clarity. The oxygen (red), nitrogen (dark blue) and sulfur atoms (yellow) are presented in the stick structure. The Mg^++^ (green) and Zn^++^ (indigo blue) ions are represented as spheres.

**Fig. S-3.**
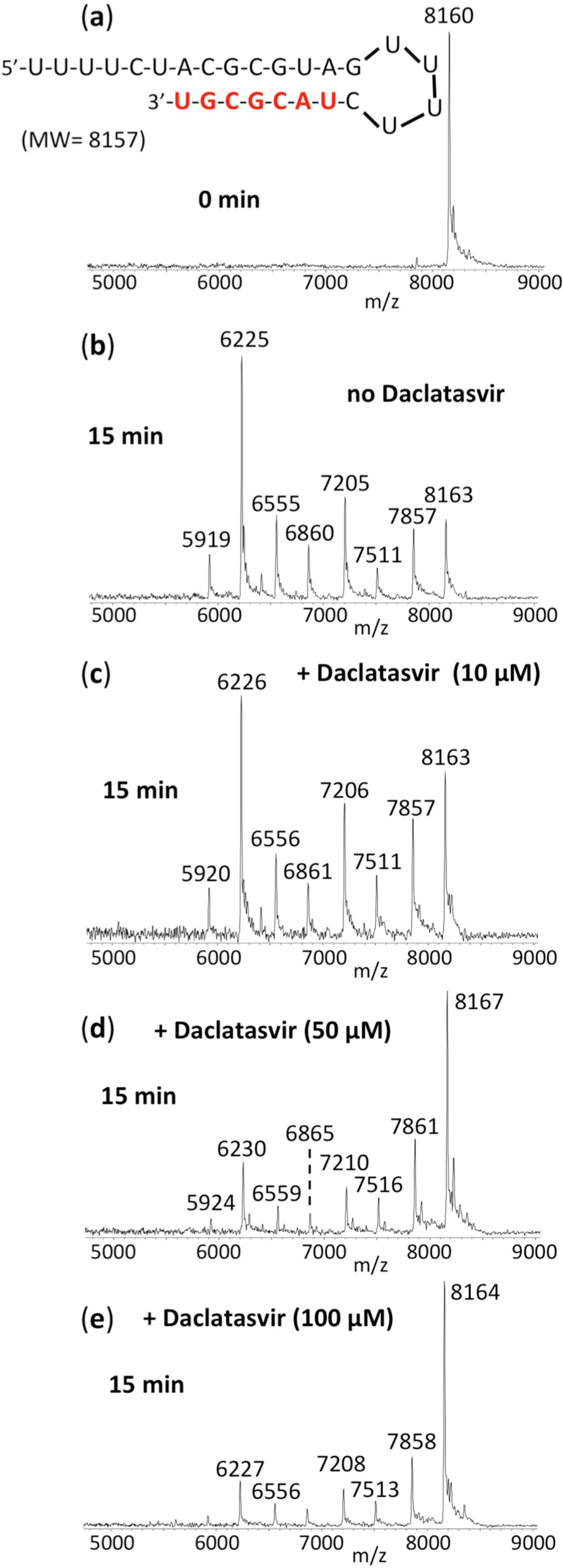
Inhibition of SARS-CoV-2 exonuclease activity by Daclatasvir. A mixture of 500 nM RNA (sequence shown at the top of the figure) and 50 nM SARS-CoV-2 pre-assembled exonuclease complex (nsp14/nsp10) was incubated in buffer solution at 37 °C for 15 min in the absence (b) and presence of varying amounts of Daclatasvir dihydrochloride (c-e). In this set of experiments, the water-soluble dihydrochloride form of Daclatasvir was used without DMSO. The RNA (a) and the products of the exonuclease reaction (b-e) were analyzed by MALDI-TOF MS. The signal intensity was normalized to the highest peak. The peak at 8160 Da corresponds to the intact RNA (8157 Da expected). In the absence of Daclatasvir, exonuclease activity caused nucleotide cleavage from the 3’-end of the RNA as shown by the 7 lower molecular weight fragments corresponding to cleavage of 1-7 nucleotides (b). With increasing amounts of Daclatasvir, exonuclease activity was reduced as shown by the reduced intensities of the fragmentation peaks and increased intact RNA peak (c-e).

**Fig. S-4:**
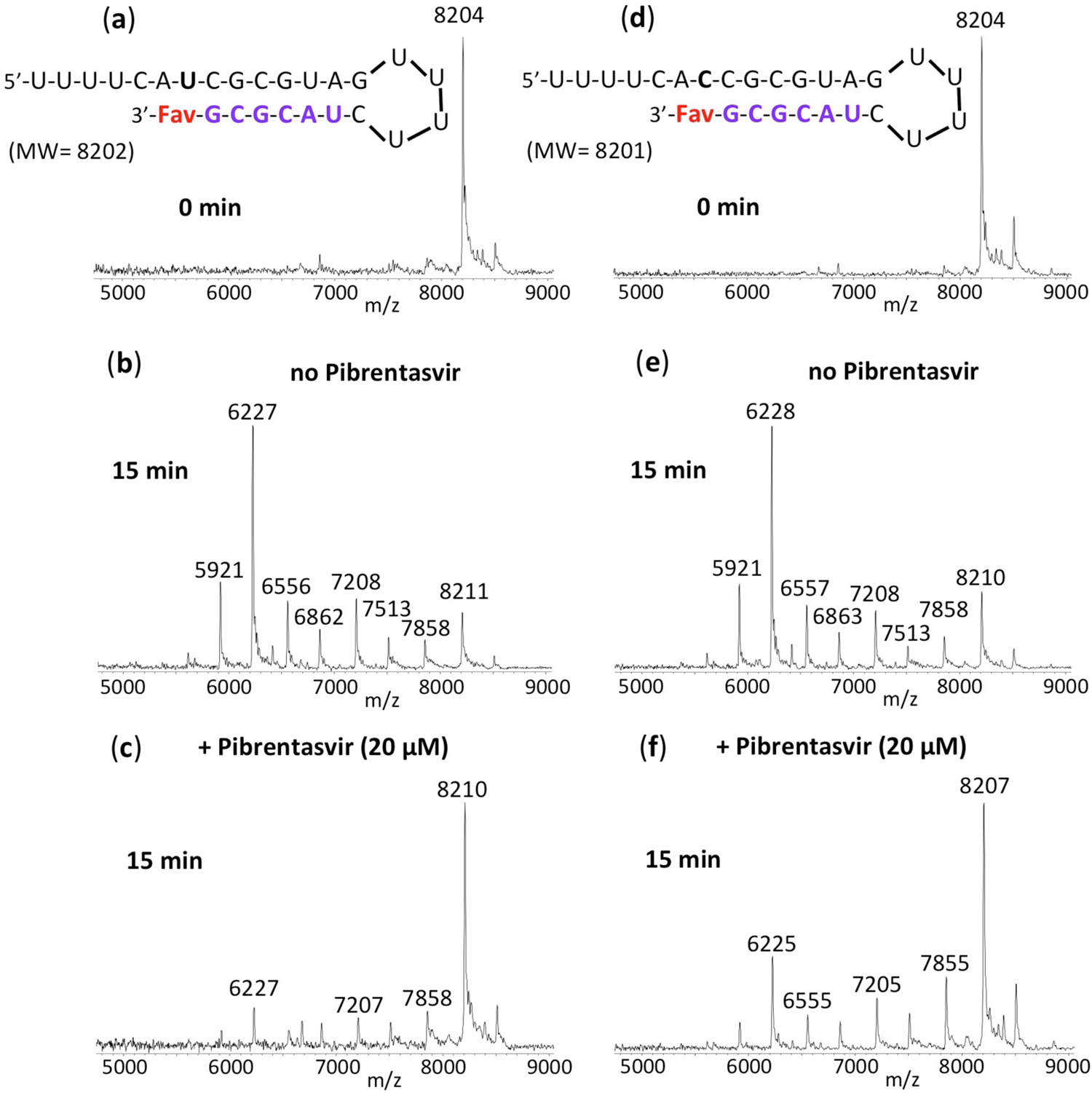
Inhibition of SARS-CoV-2 exonuclease activity by Pibrentasvir for Favipiravir (Fav) terminated RNAs. A mixture of 500 nM RNAs (sequences shown at the top of the figure) and 50 nM SARS-CoV-2 pre-assembled exonuclease complex (nsp14/nsp10) were incubated in buffer solution at 37 °C for 15 min in the absence (b, e) and presence of 20 µM Pibrentasvir (c, f). The intact RNAs (a, d) and the products of the exonuclease reactions (b, c, e, f) were analyzed by MALDI-TOF MS. The signal intensity was normalized to the highest peak. In the absence of Pibrentasvir, exonuclease activity caused nucleotide cleavage from the 3’-end of the RNA as shown by the lower molecular weight fragments corresponding to cleavage of 1-7 nucleotides (b, e). When 20 µM Pibrentasvir was added, exonuclease activity was reduced as shown by the reduced intensities of the fragmentation peaks and increased peak height of the intact RNA (c, f). These results indicate that the SARS-CoV-2 exonuclease activity is substantially inhibited by Pibrentasvir for Favipiravir terminated RNAs.

**Fig. S-5:**
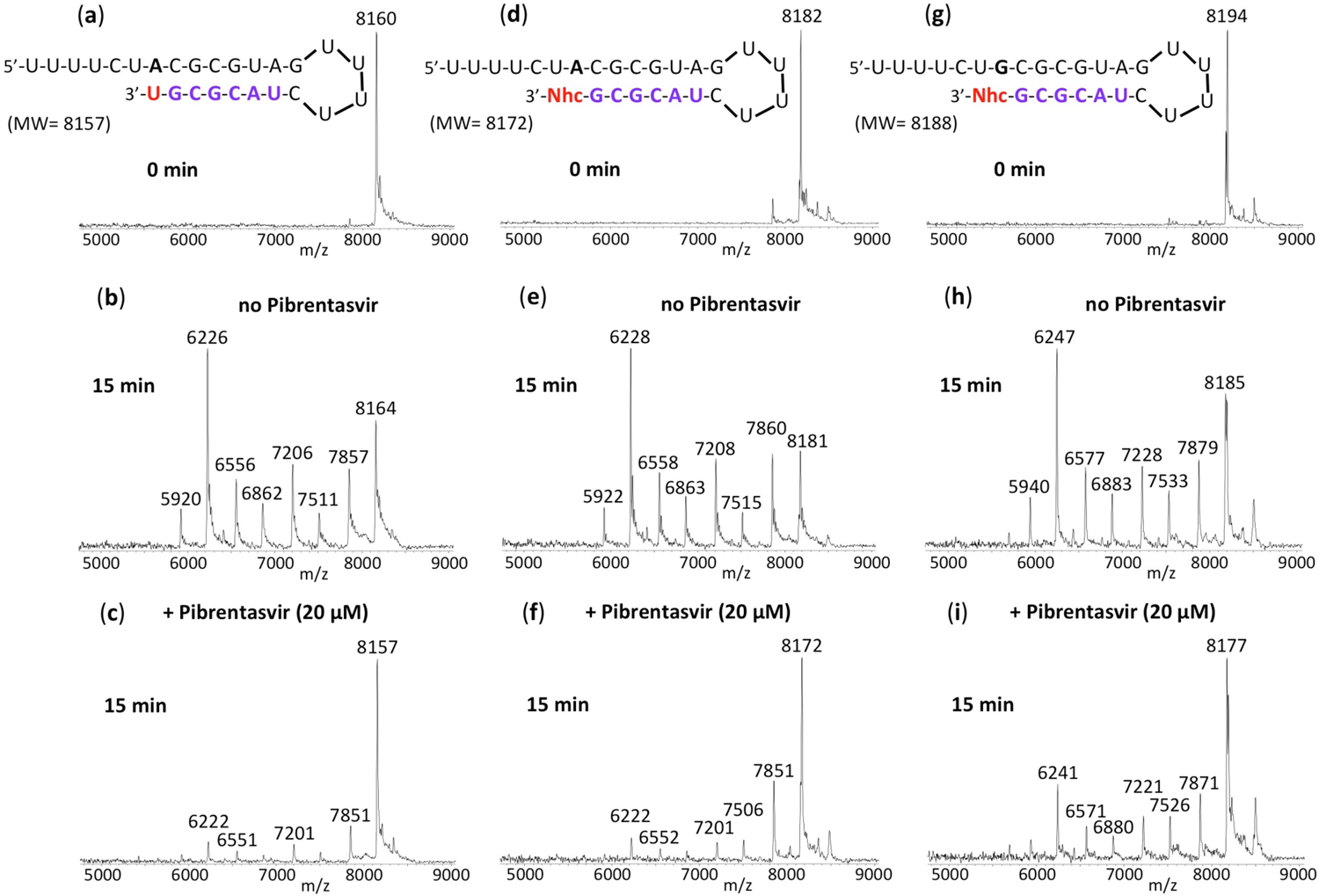
Inhibition of SARS-CoV-2 exonuclease activity by Pibrentasvir for natural RNA and N^4^-hydroxycytidine (Nhc) terminated RNAs. A mixture of 500 nM RNAs (sequences shown at the top of the figure) and 50 nM SARS-CoV-2 pre-assembled exonuclease complex (nsp14/nsp10) were incubated in buffer solution at 37 °C for 15 min in the absence (b, e, h) and presence of 20 µM Pibrentasvir (c, f, i). The intact RNAs (a, d, g) and the products of the exonuclease reactions (b, c, e, f, h, i) were analyzed by MALDI-TOF MS. The signal intensity was normalized to the highest peak. In the absence of Pibrentasvir, exonuclease activity caused nucleotide cleavage from the 3’-end of the RNA as shown by the lower molecular weight fragments corresponding to cleavage of 1-7 nucleotides (b, e, h). When 20 µM Pibrentasvir was added, exonuclease activity was reduced as shown by the reduced intensities of the fragmentation peaks and increased peak height of the intact RNA (c, f, i). These results indicate that the SARS-CoV-2 exonuclease activity is substantially inhibited by Pibrentasvir for NHC terminated RNAs.

**Fig. S-6:**
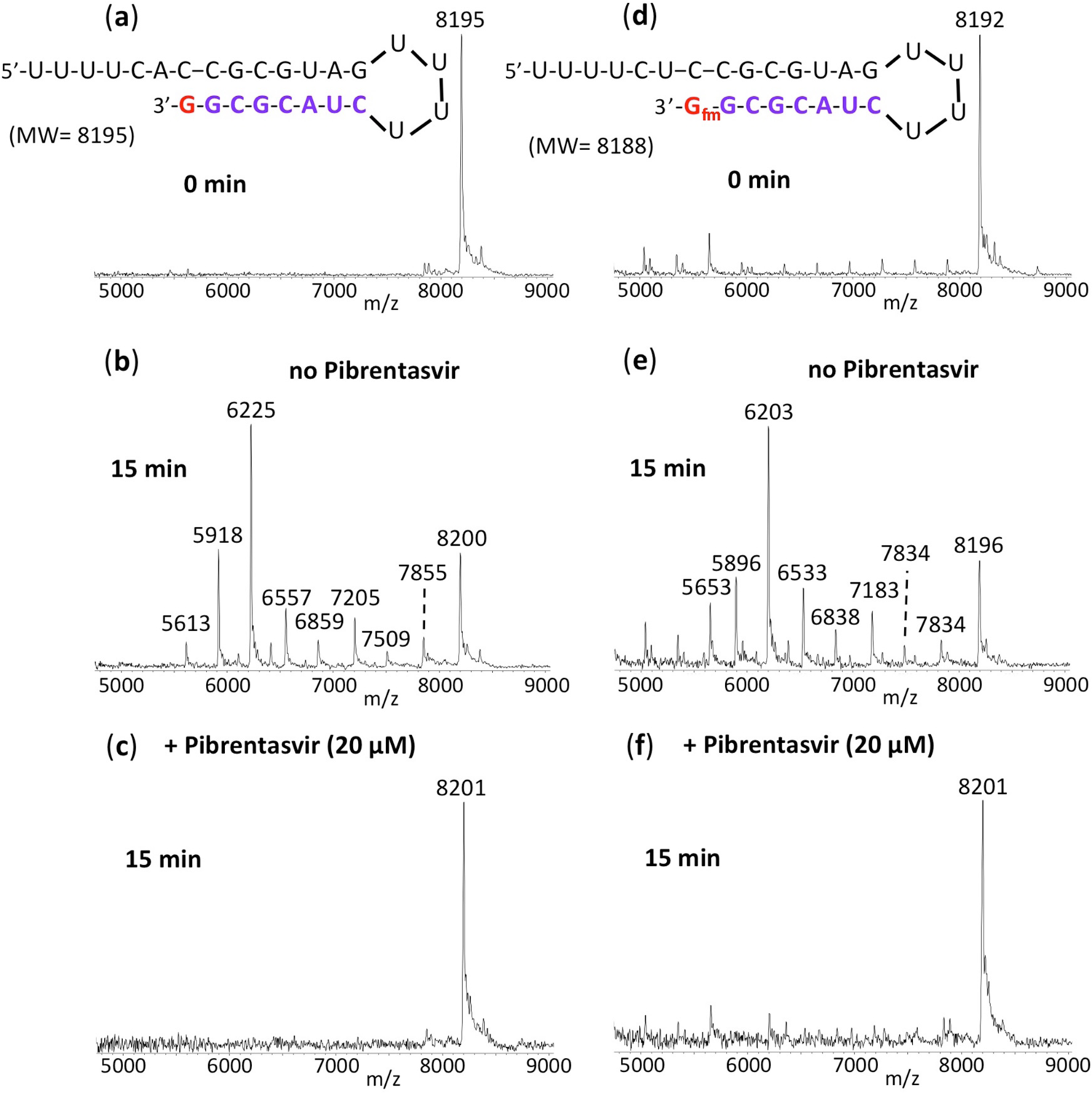
Inhibition of SARS-CoV-2 exonuclease activity by Pibrentasvir for natural RNA and AT-9010 (G_fm_) terminated RNA. A mixture of 500 nM RNAs (sequences shown at the top of the figure) and 50 nM SARS-CoV-2 pre-assembled exonuclease complex (nsp14/nsp10) were incubated in buffer solution at 37 °C for 15 min in the absence (b, natural RNA; e, G_fm_ terminated RNA) and presence of 20 µM Pibrentasvir (c, natural RNA; f, G_fm_ terminated RNA). The intact RNAs (a, d) and the products of the exonuclease reactions (b-f) were analyzed by MALDI-TOF MS. The signal intensity was normalized to the highest peak. In the absence of Pibrentasvir, exonuclease activity caused nucleotide cleavage from the 3’-end of the natural RNA as shown by the lower molecular weight fragments corresponding to cleavage of 1-8 nucleotides (b, e). When 20 µM Pibrentasvir was added, exonuclease activity was almost completely abolished as shown by the near absence of the fragmentation peaks and increased peak height of the intact RNA (c, f). These results indicate that the SARS-CoV-2 exonuclease activity is substantially inhibited by Pibrentasvir for both natural and AT-9010 (G_fm_) terminated RNA.

**Fig. S-7.**
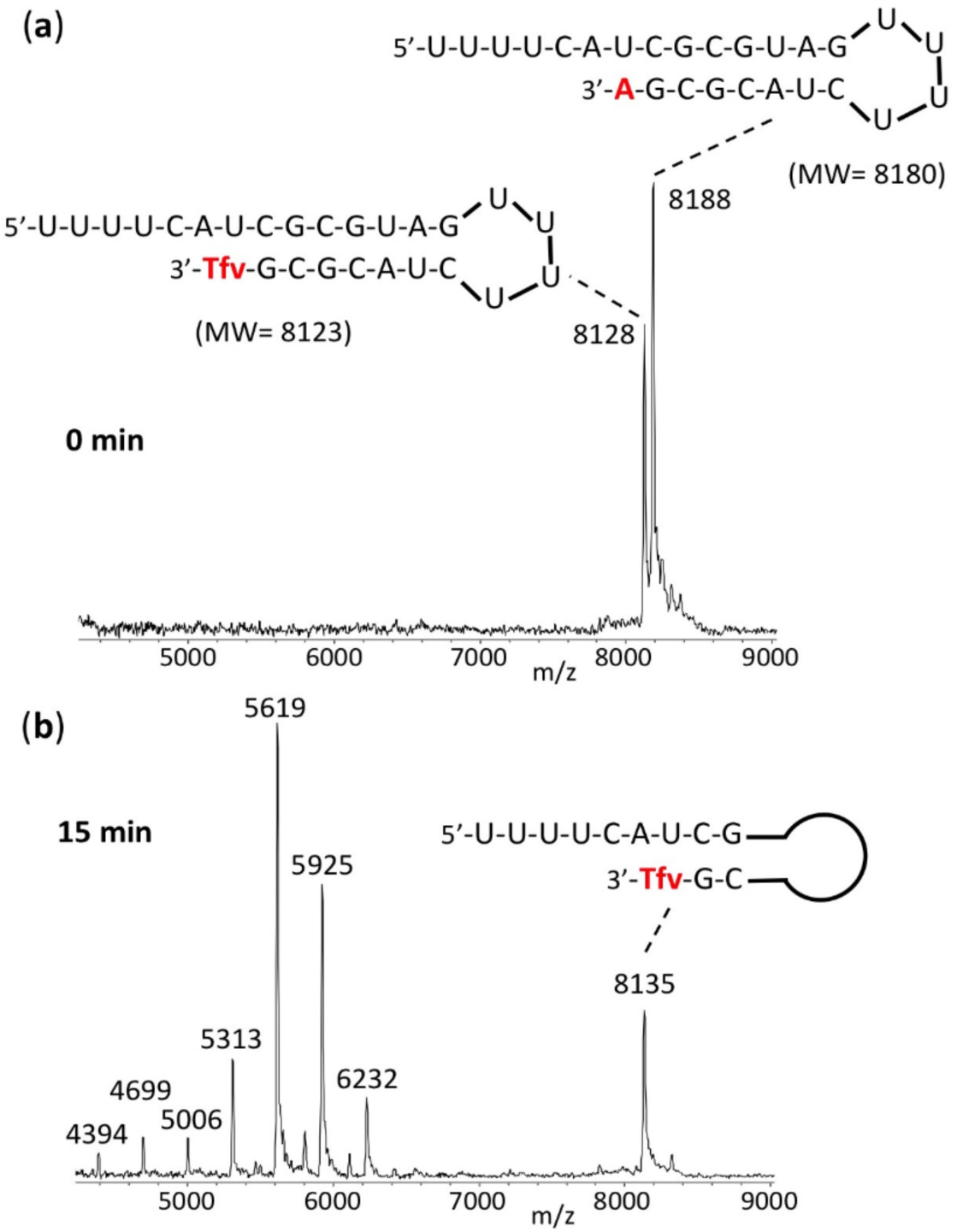
SARS-CoV-2 exonuclease activity for a mixture of natural RNA and Tenofovir (Tfv) terminated RNA. A mixture of ∼500 nM RNAs (sequences shown at the top of the figure) and 50 nM SARS-CoV-2 pre-assembled exonuclease complex (nsp14/nsp10) were incubated in buffer solution at 37 °C for 15 min (b). The untreated RNA mixture (0 min) (a) and the products of the exonuclease reactions (b) were analyzed by MALDI-TOF MS. Exonuclease activity caused nucleotide cleavage from the 3’-end of the natural RNA as shown by the lower molecular weight fragments corresponding to cleavage of up to 12 nucleotides (b). No detectable MS peak was observed for the full-length natural RNA. However, a large MS peak for intact Tfv terminated RNA was observed at 8135 Da (b) demonstrating that Tfv terminated RNA shows high resistance to SARS-CoV-2 exonuclease activity compared to natural RNA.

**Fig. S-8.**
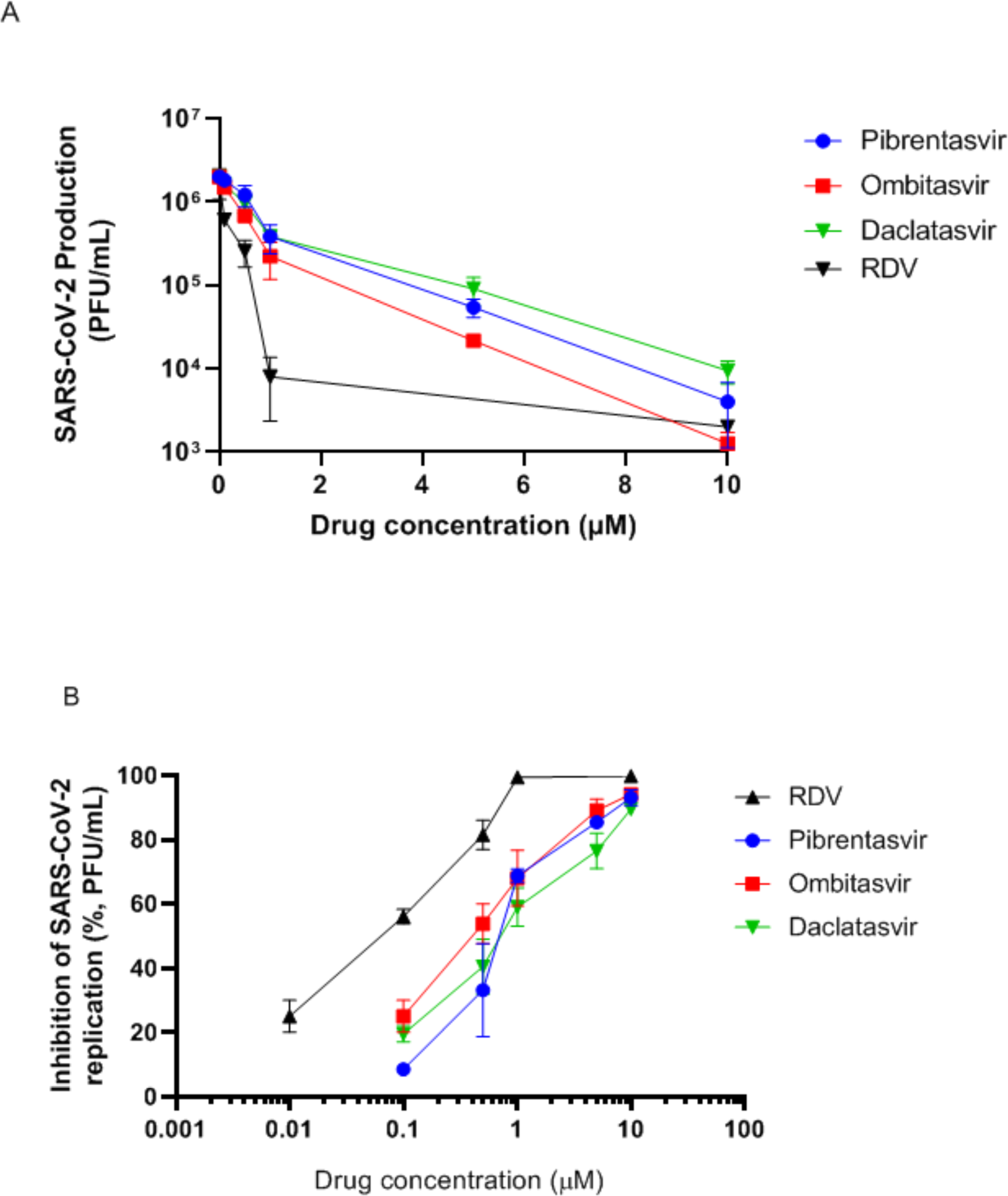
Antiviral activity of Pibrentasvir, Ombitasvir and Daclatasvir against SARS-CoV-2. Calu-3 cells, at a density of 5 × 10^5^ cells/well in 48-well plates, were infected with SARS-CoV-2 at a MOI of 0.1, for 1 h at 37 °C. An inoculum was removed and cells were washed and incubated with fresh DMEM containing 2% FBS and the indicated concentration of the drugs, including Remdesivir (RDV) as the control. Supernatants were assessed after 48-72 h. Viral replication in the culture supernatant was measured as PFU/mL by titering in Vero E6 cells. Results are displayed as virus titers (A) and percentage of inhibition (B). The data represent means ± SEM of three independent experiments.

**Fig. S-9.**
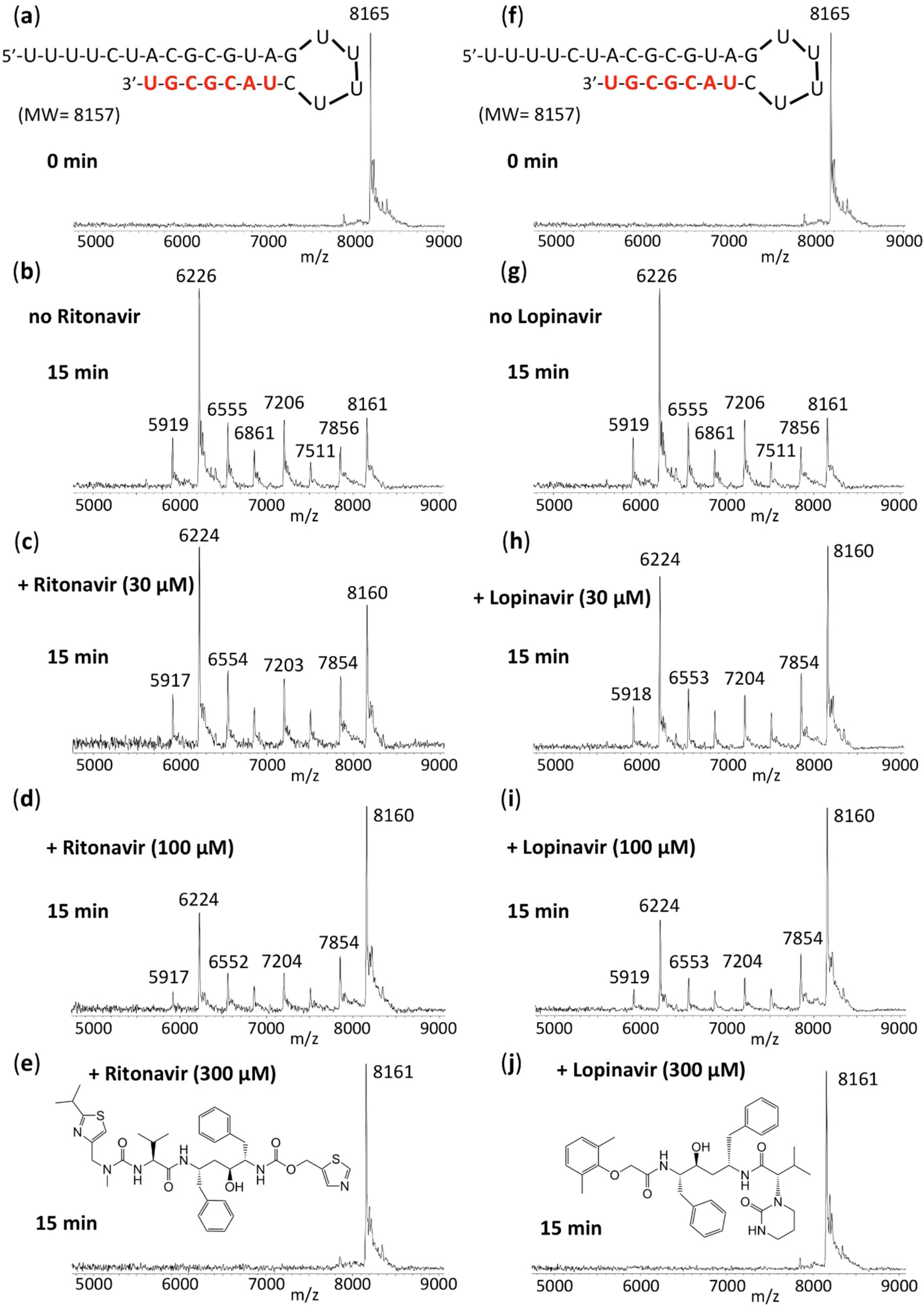
Inhibition of SARS-CoV-2 exonuclease activity by Ritonavir and Lopinavir. A mixture of 500 nM RNA (sequence shown at the top of the figure) and 30 nM SARS-CoV-2 pre-assembled exonuclease complex (nsp14/nsp10) was incubated in buffer solution at 37 °C for 15 min in the absence (b, g) and presence of 30 µM (c, h), 100 µM (d, i) or 300 µM (e, j) Ritonavir or Lopinavir (structures shown at the bottom of the figure). The RNA (a, f) and the products of the exonuclease reactions (b-e, g-j) were analyzed by MALDI-TOF MS. The signal intensity was normalized to the highest peak. The peak at 8165 Da corresponds to the intact RNA (8157 Da expected). In the absence of the NS5A inhibitors, exonuclease activity caused nucleotide cleavage from the 3’-end of the RNA as shown by the 7 lower molecular weight fragments corresponding to cleavage of 1-7 nucleotides (b, g). In the presence of Ritonavir and Lopinavir, exonuclease activity was inhibited as shown by the reduced intensities of fragmentation peaks and increased intensity of the intact RNA peak (c-e, h-j).

**Fig. S-10.**
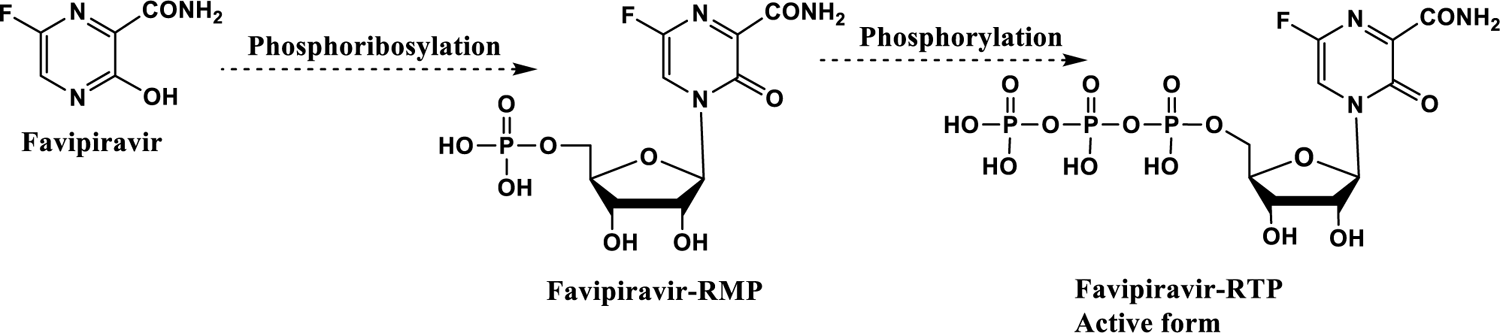
Mechanism of activation of Favipiravir. The nucleotide Favipiravir undergoes a phosphoribosylation reaction to form Favipiravir-ribofuranosyl-5’-monophosphate (RMP) followed by further phosphorylation to form the active triphosphate Favipiravir-ribofuranosyl-5’-triphosphate (RTP).

**Table S-1.** In vitro pharmacological parameters for representative viral RNA polymerase and HCV NS5A inhibitors against SARS-CoV-2 production in Calu-3 cells.

|  | EC<sub>50</sub> | CC<sub>50</sub> | SI |
| --- | --- | --- | --- |
| Drug | [µM] | [µM] |  |
| Remdesivir | 0.09 ± 0.002 | 480 ± 55 | 5333 |
| Sofosbuvir | 6.2 ± 0.3 | 512 ± 34 | 82 |
| Tenofovir (TDF) | 4.3 ± 2.1 | 91 ± 25 | 21 |
| Favipiravir | 7.8 ± 1.2 | 670 ± 118 | 85 |
| Pibrentasvir | 0.7 ± 0.2 | 22 ± 3 | 31 |
| Ombitasvir | 0.4 ± 0.05 | 70 ± 12 | 175 |
| Daclatasvir | 0.7 ± 0.08 | 38 ± 5 | 54 |

SI – Selectivity index = CC<sub>50</sub>/EC<sub>50</sub>

